# Leg choice for volitional goal-directed stepping is primarily influenced by effort rather than success: A preliminary investigation in neurotypical adults

**DOI:** 10.1101/2024.04.10.588947

**Authors:** Charalambos C. Charalambous, Eric R. Espinoza-Wade, Guilherme M. Cesar, Michaela Gerger, Yi-Hsuan Lai, Carolee J. Winstein

## Abstract

During reaching, arm choice depends on success and motor effort. Whether these factors influence leg choice for stepping behavior is unknown. Here, we conducted two experiments (1: proof-of-principle; 2: kinematic analysis) to explore whether limb selection for goal-directed stepping depends on success and/or effort under two *Choice* conditions: *Free* (choose either leg) and *Constrained* (no choice – only left leg). For both experiments, in which Free trials always preceded Constrained trials, we adapted the classic center-out target array in which right-leg dominant neurotypical adults stood in the middle of the array and stepped to pre-cued targets as accurately as possible. The *Free* condition reflected the preferred limb choice. We compared success, effort, and self-perceived difficulty between *Free* and *Constrained* trials, separately for three (Experiment 1) and two (Experiment 2) regions. Overall, in *Free* condition, participants uniformly selected the limb ipsilateral to lateral left and right targets and with slight leg dominance-based bias for central targets. Success (step accuracy and consistency/precision) did not depend on Choice condition, rather, performance improved over repeated trials. Effort (peak vertical foot lift and step path ratio) depended on Choice condition. Finally, independent of Choice condition, participants perceived posterior targets (particularly far targets) as the most difficult during non-dominant left steps. Present findings suggest that effort may influence leg choice to a greater degree than success for goal-directed stepping. Future work that probes these findings’ robustness in patients with unilateral paresis (intrinsic constraint) may advance our understanding of the motor decision processes for goal-directed mobility behaviors.

## Introduction

Recent accounts of sensorimotor control of discrete goal-directed actions have relied on a multilevel framework that progresses from action selection to movement planning to movement execution (Gallivan et al. 2018; Kim et al. 2021). Considerable work in this multilevel process has been informed by ideas from motor neuroscience (Shadmehr et al. 2016; Summerside et al. 2018) and the neuroeconomics of motor decision-making (Trommershäuser et al. 2008), with a specific focus on skilled arm aiming/reaching movements (Kim et al. 2021). Though there is considerable understanding of movement execution and planning, few studies have explicitly investigated action selection (i.e., motor decision-making) during lower extremity task performance (e.g. walking; (Dominguez-Zamora and Marigold 2021), especially for lower extremity discrete goal-directed movements. Given that part of action selection is to select a to-be-moved limb (i.e., “decide-then-act”; (Cos et al. 2011), limb selection is a form of motor decision-making. It is a binary choice, either left or right arm or leg for reaching and stepping, respectively, in humans (Wolpert and Landy 2012).

During reaching movements, humans select which arm to use based on intrinsic (e.g., handedness) and extrinsic (e.g., task context) factors (Mamolo et al. 2006; Hirayama et al. 2022). There is extensive evidence that humans typically choose the arm ipsilateral to the target (e.g., using the right arm for reaching out to a cup of coffee located on the right front); a phenomenon called hemispace/hemispheric bias (Helbig and Gabbard 2004; Bryden and Roy 2006; Coelho et al. 2013). For targets located in the anterior midline of the body and compound tasks such as reach-to-grasp, humans are biased towards selecting and then using the dominant hand (Calvert and Bishop 1998; Mamolo et al. 2006; Coelho et al. 2013; Han et al. 2013; Przybyla et al. 2013). Furthermore, “action based” models suggest that both expected success (i.e., motor-related reward, such as capturing the target with accurate and precise actions) and expected effort (i.e., motor-related biomechanical cost, a straight arm path to reduce motor effort) influence arm choice in both neurotypical adults (Cos et al. 2011; Cisek 2012; Schweighofer et al. 2015; Shadmehr et al. 2016; Wang et al. 2021) and neurologically impaired adults (Kim et al. 2018; Kim et al. 2022; Nguyen et al. 2023).

Like reaching actions, stepping can also be a discrete motor action (e.g., quick stepping backwards after an anteroposterior perturbation like chest push). In contrast to reaching, goal-directed stepping has two distinct traits. First, stepping with one leg depends on the stability of the standing limb (Day and Bancroft 2018), a factor not crucially relevant to unimanual reaching movements unless the object requires bimanual coordination. Therefore, leg selection for stepping actions may also depend on the capacity of the contralateral standing leg (e.g., no pain, sufficient strength and control) to maintain and ensure postural stability and body balance during stepping (Peters 1988). Second, unlike reaching, stepping actions can be executed either towards the front (i.e., forward stepping) or towards the back (i.e., backward stepping). Though these stepping modes share similar movement patterns, there are important sensorimotor control differences. Forward stepping uses the inputs from the visual system, whereas backward stepping has limited use of visual inputs (i.e., memory guided stepping) and may rely more on proprioceptive cues (Thomas and Fast 2000). Also, there is a functional distinction in cortical activation between the two stepping modes (Berchicci et al. 2020). Though both stepping modes provoke cortical activation in motor and premotor areas; forward stepping also provokes cortical activation in parietal areas (sensorimotor transformation) whereas backward stepping provokes cortical activation in prefrontal areas (executive control) (Berchicci et al. 2020). Furthermore, forward stepping has high motor familiarity due to walking (use of vision and forward foot pointing) whereas backward stepping has lower motor familiarity and is used mainly during protective stepping responses (Lee et al. 2014). Despite these differences from reaching, we reasoned that limb choice during volitional goal-directed stepping, which differ from compensatory stepping (McLlory and Maki 1996), might be influenced by the same intrinsic (e.g., limb dominance) and extrinsic (e.g., task context) factors that influence limb choice during reaching.

Therefore, motivated by our recent work in arm choice for reaching actions (Han et al. 2013; Schweighofer et al. 2015; Kim et al. 2022), we designed a set of experiments to investigate lower limb selection processes for volitional goal-directed stepping actions using behavioral and kinematic measures. Each experiment consisted of a single session; we adapted the classic center-out target array in which participants had to step accurately onto a pre-cued target without time restrictions. There were two Choice conditions. In the first condition, *Free*, we did not impose instructions about which limb to use, but instead instructed the participant to select the limb of their choice to step onto the pre-cued target. In the second condition, *Constrained*, there was no choice (extrinsic constraint) as we instructed the participant to use only their non-dominant limb to step onto the pre-cued target. The latter condition is motivated by the extensive work in participants with unilateral hemiparesis (i.e., intrinsic constraint) for which untapped capability is revealed when the less-affected limb (i.e., preferred limb for the action) is constrained from reaching (Sterr et al. 2014; Kim et al. 2022).

Based on prior arm reaching work, we formulated three hypotheses. First, in *Free*, the ipsilateral leg would be selected for targets located on the same side (i.e., hemispace/hemispheric bias), and a bias for the dominant leg would be observed for midline targets (i.e., dominant side bias). Second, in *Free*, the success measures would be relatively high (i.e., accurate and consistent/precise stepping) while the motor effort measures would be low (i.e., minimal foot lifts and straight foot path trajectories), particularly for ipsilateral targets. Third, in *Constrained*, when compared to *Free*, the success measures would diminish (i.e., inaccurate and inconsistent/imprecise stepping) while the effort measures would rise (i.e., higher foot lifts and curved foot path trajectories), particularly for contralateral targets.

## Experiment 1

Experiment 1 was designed as a proof-of-principle, to replicate or not, the limb selection findings reported previously in upper limb reaching studies and to compare measures of success (stepping error) of stepping actions performed under the two limb-choice conditions. Effort was not assessed directly in this experiment, but instead we captured self-reported measures to quantify perceived difficulty in which high self-perceived target difficulty may reflect perception of low success and high motor effort.

## Methods

### Participants

Twelve neurotypical and experiment-naïve students from the Division of Biokinesiology and Physical Therapy at the University of Southern California volunteered to participate in this cross-sectional study. Participants were included if they were free of musculoskeletal injuries and neurological conditions, between 20-40 years of age, and had normal or corrected to normal vision. They were excluded if they had any serious surgeries within the last 6 months or standing balance impairment. All participants read and signed a written informed consent before the experimental procedures, which was approved by the Institutional Review Board of the University of Southern California and adhered to the Declaration of Helsinki (APP-10-05784). Testing was conducted at the Motor Behavior and Neurorehabilitation Lab located at the Division of Biokinesiology and Physical Therapy.

### Experimental Procedures

All participants attended a single session which lasted less than an hour **(Fig. 1a)** and wore comfortable, flat, sport shoes. Prior to targeted stepping, baseline screening was conducted to determine leg dominance and standing balance ability (Duncan et al. 1990; Hoffman et al. 1998) **(Fig. 1a-I)**. Briefly, we used the ball kick and step-up tests to determine leg dominance; 3 trials per test. All participants chose the right leg for both tests; hence, all 12 participants were characterized as right-leg dominant. To assess balance capacity, we used the functional reaching (three trials) and single-leg stance (30 sec with eyes open) tests; all participants successfully completed both tasks with no balance deficits.

**Fig. 1.**
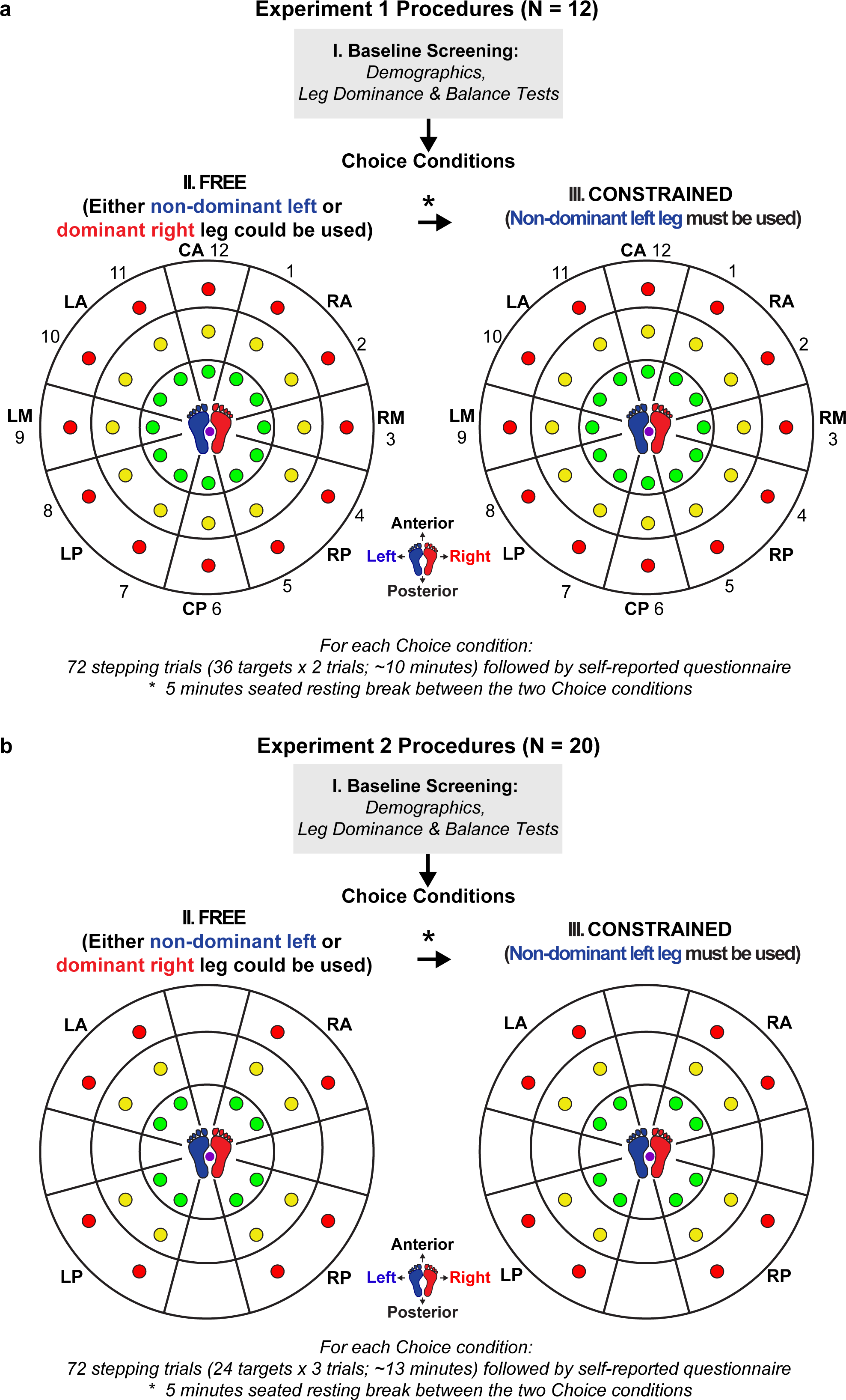
Schematic of the experimental procedures and target array used in Experiment 1 **(a)** and 2 **(b). (a)** For Experiment 1, twelve neurotypical participants attended a single session in which they were screened (I), stepped onto targets under two Choice conditions, *Free* (II) and *Constrained* (III), and completed a self-reported survey about self-perceived target difficulty after each condition. On the floor was a target clock-face array with thirty-six targets and a central start/home position (see the two-colored footprints) that was used in each condition. Targets were arranged in twelve target directions and grouped in eight target regions (clockwise numeric values): Right-Anterior (RA; 1 and 2), Right-Medial (RM; 3); Right-Posterior (RP; 4 and 5), Central-Posterior (CP; 6), Left-Posterior (LP; 7 and 8), Left-Medial (LM; 9), Left-Anterior (LA; 10 and 11), Central-Anterior (CA; 12) (See Supplementary Material for further details on the conversion of the target directions to target regions). **(b)** For Experiment 2, a new cohort of neurotypical participants also attended a single session in which they were screened (I), stepped onto targets under the same two Choice conditions, *Free* (II) and *Constrained* (III), as in Experiment 1, and completed a self-report survey about self-perceived task-difficulty after each condition. On a wooden platform was a target clock-face array with twenty-four targets and a central start/home position (see the two-colored footprints) that was used in each condition. Targets were arranged in 4 target regions: Right-Anterior (RA), Right-Posterior (RP), Left-Posterior (LP), Left-Anterior (LA). **(a & b)** Black arrows indicate the serial order of the screening and the two conditions; the asterisk (*) indicates the 5-minute seated resting break between *Free* and *Constrained*. Purple dots indicate the home center of the start/home position and target array. The two-foot prints (blue: left leg; red: right leg) surrounding the purple dot represent the start and end position before and after targeted stepping action. In both experiments, targets were arranged in three target Extents (distance between the home center of the target array and the center of the target): Green/Near (25 cm), Yellow/Middle (43 cm), Red/Far (62 cm).

The target stepping task (two choice conditions – **Fig. 1a II&III**) was an adapted version of the classic center-out target display. Importantly, in contrast to how the center-out target display has classically been used for reaching/arm aiming studies (Georgopoulos et al. 1981; Gordon et al. 1994; Cortes et al. 2017), the participant stood in the middle of the circular target array. This allowed us to explore stepping behavior in front (forward stepping with either limb), behind (backward stepping with either limb) and around the standing limb (Constrained trials with left limb to right anterior, medial, and posterior targets). There were thirty-six paper targets (diameter: 0.05 m) taped to the floor and arranged in a clock-face array: 12 target directions (i.e., 30° between directions) arranged in a clockwise fashion and 3 target extents (distance from the home center) (**Fig. 1**). The 12 target directions were converted into 8 target regions (See Supplementary Material for further details). Target region and extent constituted the task-related manipulation (i.e., independent variable of Target Region and Target Extent); these two factors can manipulate the level of success, effort, and perceived difficulty.

Two choice conditions constituted the experimental manipulation (i.e., independent variable of *Choice*); this factor can manipulate the choice of the stepping leg. In the *Free* **(Fig. 1a II)**, participants could choose either leg to step into the target. In the *Constrained* **(Fig. 1a III)**, participants had no choice (Han et al. 2013), instead they were extrinsically constrained to use the left non-dominant leg for all target regions and extents. In both conditions, participants stood with both feet at a comfortable distance apart inside a central home position and stepped when verbally cued to one of 36 targets in a pseudorandom order with no single target presented consecutively. Given that the cohort was young, neurotypical, and had no balance deficits, no overhead harness was used. Because arm motion may affect stepping limb selection (Peters 1988), we instructed participants to fold their arms across their chest when stepping. The 36-target sequence was repeated twice (72 trials per condition). The experimenter verbally indicated the designated target (“Ready, Twelve, Red, Go”) in a paced order within approximately 5 seconds (i.e., intertrial interval). Before each Choice condition, three instructions were given; the first two were the same for both conditions. The first instruction was to visually locate the target prior to stepping; but during stepping, participants were directed to gaze at a colored poster taped to the wall located 2 meters in front of them at roughly eye level. This assured that the stepping behavior was performed to a remembered visual target (i.e., memory guided stepping) for all target regions and extents. The second instruction was to distribute body weight equally on both legs when hearing the “Ready” signal, then to step onto the instructed target as accurately as possible after hearing the “Go” signal. Note, there was no mention of time to react or to move. Further, they were asked to shift body weight onto the target once the ground was contacted, and then to immediately step back to the home position. Given that we were not interested in manipulating the decision process itself (e.g., limiting time to process), we neither constrained movement time nor recorded reaction time. Participants were instructed not to correct an error trial after they had stepped. The third instruction differed between the two *Choice* conditions. For the *Free* stepping trials, participants could choose either leg for stepping whereas for the *Constrained* stepping trials, participants were instructed to step into each target using only the left leg (i.e., non-dominant leg). To ensure that participants understood the instructions and procedures for each condition, the experimenter demonstrated, and participants practiced a few trials (∼ 5) of each condition. To minimize context effects, we blocked the two conditions and put the easier one first; the *Free* trials were always administered before the *Constrained* trials. There was a five-minute rest period between conditions in which participants sat on a chair (see asterisk in **Fig 1a**), to minimize any effects of fatigue. No explicit augmented feedback was provided. All stepping trials were videotaped using a camcorder (60 Hz) and used off-line to verify limb selection and success (accuracy).

After each *Choice* condition, we administered a single item questionnaire where participants identified by circling the most difficult target(s) on a target-labeled figure. Participants were asked “Which target(s) did you find to be the most difficult?” Participants had to select at least one target and could select more than one target if desired.

## Data Analyses

By reviewing data collection videotape offline, in *Free*, the visual inspection was used to determine the limb used for stepping to each target whereas in *Constrained*, visual inspection was used to confirm that the left leg was used as instructed. We calculated three measures: the probability of choosing the right limb (proxy of limb choice), stepping error (proxy of accuracy), and perceived target difficulty (proxy of perceived low success and high motor effort). The first measure was calculated in Free only whereas the latter two were calculated in both *Free* and *Constrained* conditions. For the probability of choosing the right limb and stepping error, which both were continuous with a range 0-1, we analyzed first each target (e.g., if the values for the first and second trials were 0 and 1, then the value used for that target was 0.5) and then averaged target region for each target extent (e.g. Near RA targets: Green 1 and 2) per participant, and then we averaged across participants. For the perceived target difficulty, which was also continuous with a range 0-1, we calculated the perceived target-difficulty for each target for each participant, and then we calculated the average across all participants by target. For further details see Supplementary Material.

## Statistical Analyses

To test whether neurotypical adults select a preferred stepping limb for a given target, we ran a separate 2-way rmANOVA using 8 Target Regions x 3 Target Extents. For limb choice probability that applied in only *Free*, we anticipated that participants would choose and use the right leg always for right region targets, mainly for the central region targets and barely if at all for the left region targets. Hence, we expected that the probability of choosing the right limb would be influenced primarily by Target Region and not by Target Extent.

To test our expectations for stepping error and self-perceived task-difficulty, we ran three separate 3-way rmANOVA to determine the interactive effect of Choice × Target Region × Target Extent on left target regions (LP, LM, LA), right target regions (RA, RM, RP), and central target regions (CA, CP). For all three rmANOVAs, any interaction or main effect that included Choice would reflect the nature of limb selection preferences, whereas any effect or interaction of Target Region and Extent alone would provide statistical evidence about the effect of target location, independent of limb choice. Given the study aim and our main manipulated variable (i.e., Choice), we focused on and reported the interactions and main effects of Choice (i.e., Choice × Target Region × Target Extent, Choice × Target Region, Choice × Target Extent, Choice). To determine the locus of any significant interaction or main effect of Choice, we visually examined each measure for apparent patterns that differed for the two Choice conditions, and then determined which post-hoc tests to use with the appropriate Bonferroni correction.

Importantly, for steps into left targets, we expected that Choice would have neither a significant effect nor significant interaction on either stepping errors or self-perceived target-difficulty because these targets are ipsilateral to the stepping limb in both *Free* (i.e., it was expected that participants would select the ipsilateral left leg) and *Constrained* (i.e., participants were extrinsically constrained to use the ipsilateral left leg). However, for stepping actions into the right targets (i.e., ipsilateral right leg and contralateral left leg in *Free* and *Constrained*, respectively), we expected that Choice Condition would have a significant effect and/or interaction on both stepping error and self-perceived target-difficulty. Given that the dominant right leg would likely be chosen more often than the left for the central anteroposterior targets, we expected that Choice Condition would have neither a significant effect nor significant interaction on either stepping error or self-perceived target-difficulty.

To provide further confirmatory empirical support pertaining to limb selection processes, we tested the association between stepping error and self-perceived target-difficulty using correlation analyses separately for *Free* and *Constrained.* Given the relatively small N for targets selected by participants to be difficult, (Free = 9/36; Constrained = 11/36), we used a non-parametric test and reported Spearman’s rank correlation coefficient *(rho)*. We reasoned that if there was a significant correlation it should be a positive one in which as stepping error increases, self-perceived target difficulty would also increase.

The statistical analyses used are summarized in Table S1 (See Supplementary Material). We present all data as mean ± 1SD and used SPSS v25 (IBM Corp, Armonk, NY, USA) for all statistical analyses, with level of significance set at α = 0.05.

## Results

All twelve participants (5 men, mean ± SD, age = 30.2 ± 5.0 years) completed the stepping task in both conditions (i.e., feasibility) without any adverse events, including falls or fatigue (i.e., safety). **Fig. 2** illustrates the results using color-coded heat maps for each Choice condition and each of the three dependent measures. Visual inspection shows that the probability of choosing the right leg for right target regions (RA, RM, RP) was high (i.e., always right leg; red) and zero (never the right leg; blue) for the left target regions in *Free*, whereas both central target regions (anterior – CA and posterior – CP) were mixed with a slight bias for the right-dominant limb (i.e., right leg in more than 50% of attempts) (**Fig. 2a**). As instructed, the probability of choosing the right leg in *Constrained* was zero (never the right leg; blue) (**Fig. 2b**). Stepping error was generally low in *Free* (**Fig. 2c**), but higher in *Constrained*, especially for right medial (RM) and posterior (RP) far targets (**Fig. 2d**). Finally, self-perceived target difficulty was the highest for the central posterior (CP) targets, especially for the far central posterior region in *Free* (**Fig. 2e**). By contrast, in *Constrained*, self-perceived target difficulty was the highest in the right medial (RM) and posterior (RP) far targets (**Fig. 2f**).

**Fig. 2.**
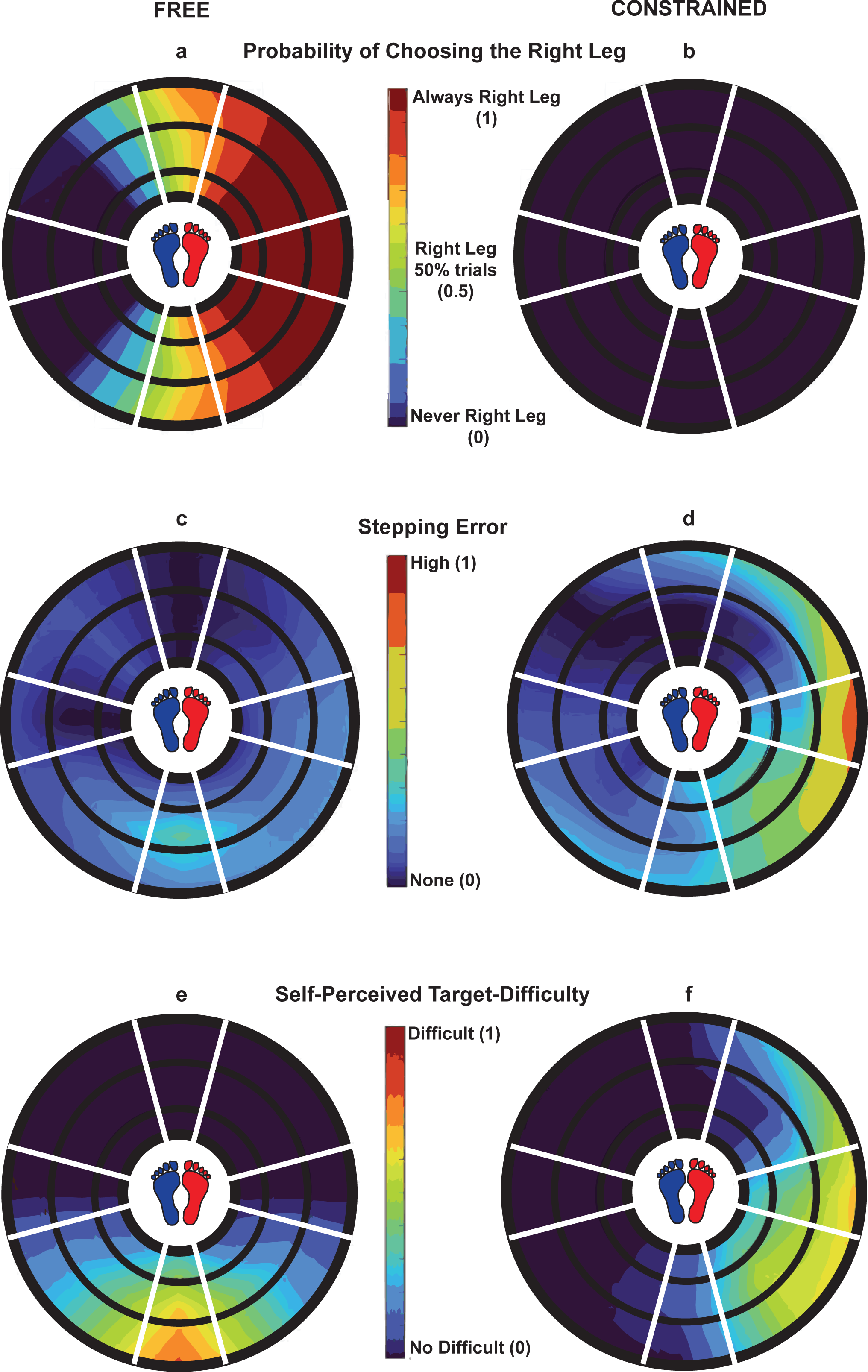
Heatmaps of the behavioral measures during *Free* (left column, **a, c, & e**) and *Constrained* (right column, **b, d, & f**) in Experiment 1. (Top row: **a & b**) Probability of choosing the right limb in *Free* (**a**) and *Constrained* (**b**). Red indicates that all subjects always selected the right leg (group mean = 1) whereas blue indicates that all subjects always selected the left leg (group mean = 0). (Middle row: **c & d**) Stepping error in *Free* (**c**) and *Constrained* (**d**). Red indicates that the stepping error was high (group mean = 1; always inaccurate) whereas blue indicates no stepping error (group mean = 0; always accurate). (Bottom row: **e & f**) Self-perceived target difficulty in *Free* (**e**) and *Constrained* (**f**). Red indicates that a target was perceived as the most difficult (mean across selected targets = 1) whereas blue indicates that a target was perceived to be not difficult (easy; mean across selected targets = 0). (**a-f**) The three colored rings in each heatmap indicate the three target extents (inner ring – near; middle ring – middle; outer ring – far)

### Stepping Limb Use per Choice Condition

In *Free*, as expected, there was an effect of Target Region on the probability of choosing the Right limb (*F* (1, 7) = 82.192, *p* = 0.001). Specifically, participants chose the limb ipsilateral to the target’s location, 100% of the time except for one participant who stepped into “Red 7” located in the LP target region for a single trial, which was the first of the two, using the right leg—thus illustrating a bias towards choosing the dominant right leg for central target regions **(Fig. 2a)**. For *Constrained*, as designed, all participants used their left leg to step onto all target regions and extents **(Fig. 2b)**.

**Table 1** summarizes the statistical results for stepping error and self-perceived target difficulty for the stepping actions into left, right, and central targets in both Choice conditions; results of any post-hoc analyses are presented in the text.

**Table 1:** Summary of the results for the repeated measures ANOVA in Experiment 1.

| DVs/IVs | Choice<br>Condition<br>×<br>Target<br>Region<br>×<br>Target<br>Extent | Choice<br>Condition<br>×<br>Target Region | Choice<br>Condition<br>×<br>Target<br>Extent | Choice<br>Condition |
| --- | --- | --- | --- | --- |
| *Stepping into the LEFT Targets in Free and Constrained* |  |  |  |  |
| Stepping<br>Error | $p=0.474$;<br>$F_{(1,4)}=0.90$ | $p=0.165$;<br>$F_{(1,2)}=1.96$ | $p=0.721$;<br>$F_{(1,2)}=0.33$ | $p=0.894$;<br>$F_{(1,1)}=0.02$ |
| Self-Perceived<br>Target<br>Difficulty | $p=0.053$;<br>$F_{(1,4)}=2.54$ | **$p=0.008$;**<br>**$F_{(1,2)}=6.06$** | $p=0.102$;<br>$F_{(1,2)}=2.54$ | **$p=0.032$;**<br>**$F_{(1,1)}=6.06$** |
| *Stepping into the RIGHT Targets in Free and Constrained* |  |  |  |  |
| Stepping<br>Error | $p=0.106$;<br>$F_{(1,4)}=2.26$ | **$p=0.033$;**<br>**$F_{(1,2)}=3.99$** | **$p=0.024$;**<br>**$F_{(1,2)}=4.42$** | **$p=0.001$;**<br>**$F_{(1,1)}=29.14$** |
| Self-Perceived<br>Target<br>Difficulty | $p=0.166$;<br>$F_{(1,4)}=1.71$ | **$p=0.001$;**<br>**$F_{(1,2)}=12.46$** | $p=0.122$;<br>$F_{(1,2)}=2.32$ | **$p=0.005$;**<br>**$F_{(1,1)}=11.99$** |
| *Stepping into the CENTRAL Targets in Free and Constrained* |  |  |  |  |
| Stepping<br>Error | $p=0.492$;<br>$F_{(1,4)}=0.74$ | $p=0.339$;<br>$F_{(1,2)}=1.00$ | $p=0.146$;<br>$F_{(1,2)}=2.12$ | $p=0.491$;<br>$F_{(1,1)}=0.51$ |
| Self-Perceived<br>Target<br>Difficulty | **$p=0.010$;**<br>**$F_{(1,4)}=5.73$** | **$p=0.001$;**<br>**$F_{(1,2)}=42.31$** | **$p=0.010$;**<br>**$F_{(1,2)}=0.73$** | **$p=0.001$;**<br>**$F_{(1,1)}=42.30$** |

DVs: dependent variables; IVs: independent variables

### Stepping into Left Target Regions

As anticipated, for the left target steps in which the left ipsilateral leg was used in both *Free* and *Constrained*, there was neither interaction nor effect of Choice for stepping error; however, there was just a significant Choice × Target Region interaction for self-perceived target-difficulty **(Table 1**, see the three left target regions in **Fig. 2c&d** and **2e&f)**. Specifically, participants perceived the left posterior targets (LP; i.e., no visual inputs of the target and backward walking) to be more difficult in *Free* than in *Constrained* (t _(11)_ = 2.462; *p* = 0.032).

### Stepping into the Right Target Regions

For the right target steps, in which right and left leg were used in accord with *Free* and *Constrained*, respectively, there were significant: 1) Choice × Target Region and Choice × Target Extent interactions for stepping error and 2) Choice × Target Region interaction for self-perceived target-difficulty **(Table 1**, see the three right target regions in **Fig. 2c&d** and **2e&f)**. Specifically, for *Constrained* (i.e., left leg was used) compared to *Free* (i.e., right leg was used), participants: 1) demonstrated greater stepping error for the medial (RM; t_(11)_ = –4.103; *p* = 0.002) and posterior (RP; t_(11)_ = –3.447; *p* = 0.005) target regions and for the far (t_(11)_ = –4.267; *p* = 0.001) target extent and 2) perceived only the right medial (RM) target regions to be more difficult (t_(11)_ = –4.304; *p* = 0.001).

### Stepping into the Central Target Regions

For the central anterior (CA) and posterior (CP) central target steps, in which either leg was used in *Free* and left leg in *Constrained*, there was neither an interaction nor effect on stepping error; however, there was a significant Choice × Target Region × Target Extent interaction for self-perceived target difficulty **(Table 1**, see the two central anteroposterior target regions in **Fig. 2c&d** and **2e&f)**. Specifically, for *Free* compared to *Constrained*, participants perceived only the middle (t _(11)_ = 3.924; *p* = 0.002) and far (t _(11)_ = 7.416; *p* < 0.001) posterior (CP) targets to be more difficult.

### Correlations

There was not a significant correlation in the *Free* condition (*rho* = 0.596, *p* = 0.090; **Fig. 3a**); however, there was a significant positive correlation between stepping error and self-perceived target-difficulty in the *Constrained* condition (*rho* = 0.704, *p* = 0.016; **Fig. 3b**).

**Fig. 3.**
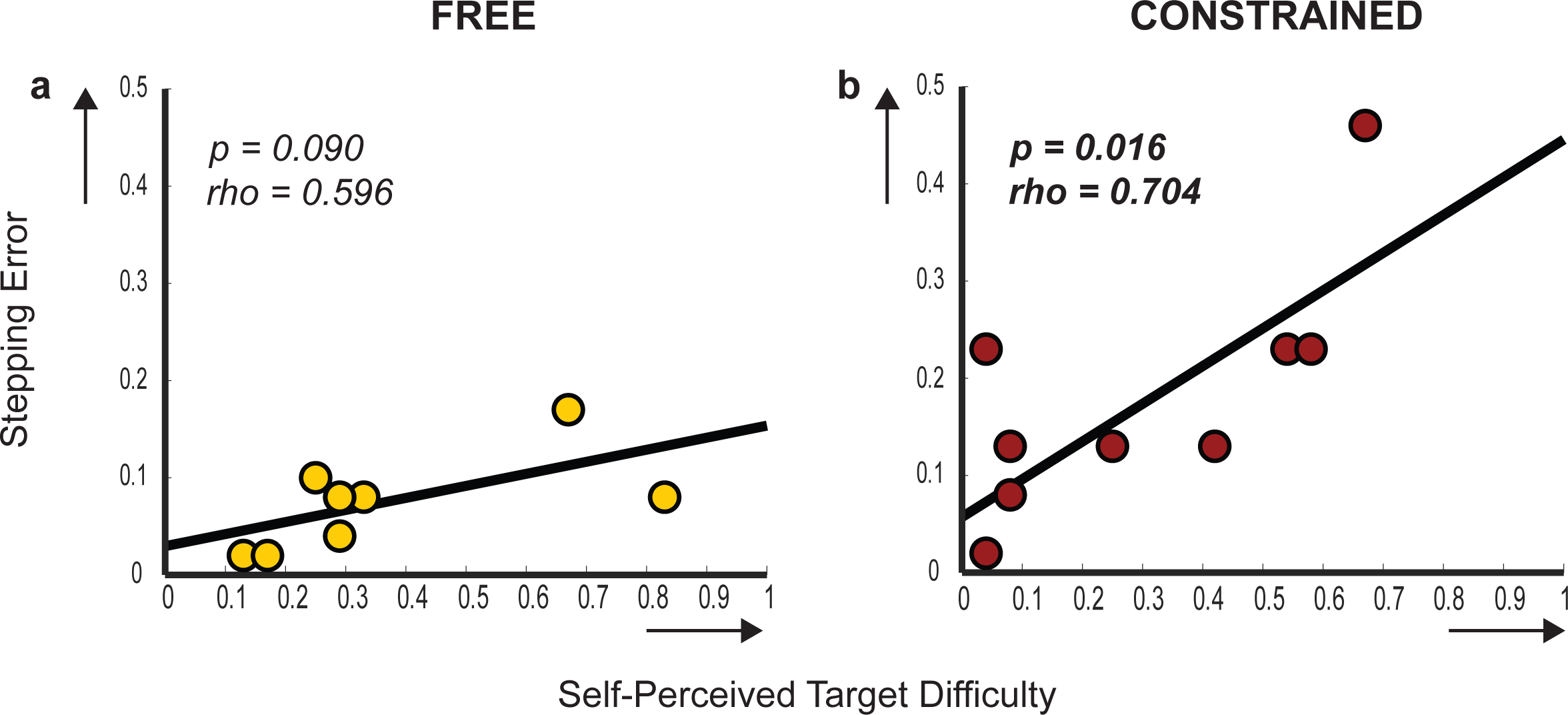
Non-parametric Spearman’s correlations between the stepping error and self-perceived target difficulty during *Free (a)* and *Constrained (b)*. In *Free* and *Constrained*, 9 and 11 data points (i.e., number of targets that were selected at least by one participant) were used, respectively; note that certain data points are superimposed (1 in *Free* and 2 in *Constrained*). The horizonal and vertical arrows in x– and y-axis indicate increase in self-perceived target difficulty and stepping error, respectively.

## Summary

Experiment 1 established the proof-of-principle, safety (i.e., no participant reported any adverse events), and feasibility (i.e., all participants completed the experimental procedures without fatigue) for the experimental paradigm and for the most part, replicated the limb selection findings reported in prior arm reaching studies. More specifically, the probability of choosing the ipsilateral limb in *Free* provides evidence that the experimental approach used here can probe limb selection processes for discrete stepping (i.e., proof-of-principle). For the left and right target regions, participants uniformly selected the leg ipsilateral to the target, clear evidence for the hemispace/hemispheric bias effect previously reported in reaching studies (Helbig and Gabbard 2004; Coelho et al. 2013).

For steps into the central anteroposterior target regions (CA and CP), participants selected either leg with a bias towards the right-dominant leg (See **Fig. 2a**). Similarly, neurotypical adults are biased towards selecting and then using the dominant hand for targets located in the anterior midline of the body (Helbig and Gabbard 2004; Coelho et al. 2013; Przybyla et al. 2013). Here, this bias towards using the dominant leg was evident for targets located both anterior and posterior to the body. The presence of this bias is consistent with the notion that right-handed adults prefer using their left leg for standing support and postural stability (Peters 1988). Though we did not record hand dominance, we suspect that most of our cohort was right-hand dominant. Further, we found that as target extent increased, participants preferred using the right-dominant leg for stepping and therefore relying on the left-non-dominant leg for postural stability. In addition, the direction of stepping into central targets is like that for typical forward and backward stepping; extending this logic, the bias for the right leg for these central targets might stem from its natural use for gait initiation and reactive backward stepping (Dessery et al. 2011).

Beyond the findings from reaching studies, the results pertaining to stepping error and self-perceived target-difficulty provide preliminary evidence that expected success (i.e., inverse of stepping error) may be important for stepping limb selection, whereas perceived difficulty (self-perceived target difficulty) may be important only under conditions of higher task demands (e.g., limited vision, less familiar and awkward conditions).

Success (i.e., stepping error) decreased (i.e., less accuracy) only for stepping into the right contralateral targets using the left leg especially for targets that were not entirely in the visual field and were located at the farthest extents. Therefore, the preferred strategy for using the ipsilateral leg relative to targets is one way to ensure more accurate stepping actions. We speculate that this may stem from the familiarity/more prior practice stepping into those directions. In addition, the preferred strategy minimizes the biomechanical challenges (i.e., no need to step around the standing limb).

Surprisingly, stepping direction, either forward or backward, had no effect on success for the central anteroposterior target regions (CA and CP); this contrasted with what was found for left and right targets. These findings suggest that these naturally used anteroposterior midline stepping actions can be successfully executed independent of limb choice and stepping direction. One possible explanation for this stand-out result is that these specific midline stepping behaviors are so well practiced, they may reflect the qualities of an especial skill (Keetch et al. 2005).

Subjective difficulty was prominent for three specific regions: the left posterior (LP) target region for which the left leg was moved backwards (i.e., no visual inputs), the central posterior (CP) for which either leg was moved backwards (i.e., no visual inputs), and the right medial (RM) target region for which the left leg was moved around the right standing limb (i.e., biomechanical challenge). These findings suggest that the lack of vision (i.e., LP and CP – targets without vision) and biomechanical challenge (i.e., RM – targets located in a position in which awkward movement is required) may influence judgements of difficulty. Based on these results, and by logical extension, it follows that stepping into awkwardly located targets with inherent biomechanical challenges and posteriorly located targets with no vision can be avoided by simply reverting to the preferred strategy (i.e., using the ipsilateral leg), if possible.

Furthermore, the strong, positive correlation between stepping error and self-perceived target-difficulty was found only for the *Constrained* condition in which we anticipated both stepping error and self-perceived target-difficulty to be increased. The results of Experiment 1 provide preliminary evidence that this extrinsically constrained condition can potentially unmask latent motor control capability even in uncommon, less preferred, and more difficult task conditions. Specifically, in *Free,* participants selected the right leg to step into the right targets to ensure success and because they did not perceive those targets to be difficult whereas in *Constrained,* those same participants demonstrated the capacity to use the left leg to step into the right targets, though with higher stepping error (i.e., limited success) and higher perceived difficulty.

Experiment 1 provides an initial proof-of-principle that our experimental paradigm could be used for the purposes intended. Experiment 2 was designed to provide quantitative evidence (i.e., kinematics of stepping behavior) to identify the role of success and/or motor effort more clearly on limb selection for the same goal-directed stepping task.

## Experiment 2

Experiment 2 was designed to replicate and extend the face validity and feasibility of the experimental method of Experiment 1. Specifically, we aimed to compare kinematically derived measures of success (constant error and variable error) and effort (peak vertical foot lift, step path ratio), and a subjective measure of perceived difficulty (target-specific perceived difficulty) from stepping actions performed under the same *Free* and *Constrained* conditions.

## Methods

### Participants

A new cohort of 20 neurotypical and experiment-naïve adults from the Division of Biokinesiology and Physical Therapy at the University of Southern California volunteered to participate. The inclusion and exclusion criteria, the informed consent process, leg dominance (i.e., again, all participants here were right-leg dominant) and balance tests, and the instructions for attire and shoes were the same as for Experiment 1. Testing was conducted at the Human Performance Laboratory located at the Competitive Athlete Training Zone (CATZ) Physical Therapy and Sports Performance facility in Pasadena, CA, USA.

## Experimental Procedures

Procedures (e.g., participant burden, baseline screening, order of conditions and target extents were the same as for Experiment 1 **(Fig. 1b)**; however, we implemented four experimental modifications motivated by the observations and results of Experiment 1 to refine the methods employed in Experiment 2 (see Supplementary Material for the detailed description of these modifications). Briefly, we eliminated both the central and medial targets; therefore, the target array consisted of 24 targets. Target extents were the same; the new, modified target array is outlined in **Fig. 1b**. Second, we included a relevant set of kinematic measures to better characterize and quantify success and effort. Third, the number of trials per target was increased from 2 to 3; total number of trials per condition was still 72. Fourth, we refined the assessment of self-perceived target difficulty. For a detailed description of the experimental procedures please see Supplementary Material.

To capture the stepping behavior (i.e., limb selection) and bilateral foot kinematics we used a camcorder (60 Hz) and 10-camera Qualisys motion analysis system (Qualisys, Gothenburg, Sweden), respectively. A single reflective marker tracked by the motion analysis system was affixed to each shoe at the level of the second metatarsal. As in Experiment 1, we captured self-perceived difficulty, at the end of each limb choice condition. However, we used a new survey question in which participants were asked *“Overall, which target did you find the most challenging”*. In contrast to Experiment 1, all participants had to choose just one target, which they perceived to be the most challenging during each Choice condition, using a target-labeled figure (See **Fig. S1** in Supplementary Material).

## Data Analyses

The stepping limb used in each Choice condition was determined by visual inspection of the reflective marker with the Qualisys Track Manager software. In *Free*, visual inspection was used to determine the limb used for stepping on each target whereas in *Constrained*, visual inspection was used to confirm that the non-dominant left leg was used for all stepping trials.

After signal processing the three-dimensional position data of the toe markers, kinematic measurers were calculated to quantify success and effort; for detailed description of data analysis see Supplementary Material. We quantified actual success (i.e., accuracy and consistency/precision) by calculating the constant error and variable error. Low constant error denotes high accuracy while small variable error denotes high consistency/precision of motor actions (Schmidt et al. 2019). We quantified motor effort by calculating the peak vertical foot lift and step path ratio. Given that stepping actions in our experiments resemble the swing phase of walking (i.e., foot off and foot contact with the same leg), we calculated foot lift because it is a major contributor to walking energy cost (i.e., the higher the foot lift the higher the energy cost) (Wu and Kuo 2016; Faraji et al. 2018). Therefore, the higher the peak vertical foot lift the higher the motor effort. Step path ratio is a unitless measure and a measure of straightness of the foot path (Stewart et al. 2013). Its minimum value is 1 which implies that the path is straight (i.e., foot path distance is equal to movement distance), effort is low and most efficient. On the other hand, any step path ratio values greater than 1 implies that the step path to the target is not straight (i.e., foot path distance is greater than movement distance) and that motor effort is higher and less efficient.

Using the answers from the self-perceived task difficulty question, we captured the frequency of the most challenging target across participants for both Choice conditions. The target regions and extents with the most challenging target would add qualitative evidence from the perspective of the performer of the degree to which target characteristics (i.e., region, extent) play a role in limb selection.

## Statistical Analyses

Now without any central targets, we expected that, during *Free* trials participants would choose the left and right leg for the left and right targets, respectively. Therefore, we ran a frequency analysis of the limb used per target region to verify the replication of Experiment 1 results.

Given our expected results for limb selection, we ran two separate 3-way rmANOVA to determine the interactive effect of Choice × Target Region × Target Extent on constant error, variable error, peak vertical foot lift, and step path ratio on left target regions (LP, LA), and on right target regions (RA, RP; See **Table S2** in Supplementary Material). As in Experiment 1, we focused on and reported any interaction or effect of Choice; we employed a similar approach for post-hoc analyses as in Experiment 1.

For stepping actions into the left targets, we expected neither interaction nor effect of Choice for any kinematic measure because these targets are ipsilateral to the stepping limb (i.e., left leg) in both *Free* and *Constrained*. We expected stepping actions with the left leg in both conditions would be accurate, consistent/precise, with relatively effortless foot lift and relatively straight foot paths. For stepping actions into right targets (i.e., ipsilateral right leg and contralateral left leg in *Free* and *Constrained*, respectively), we expected that Choice condition will have an interaction and/or effect for each of the four kinematic measures. We expected stepping actions with the left leg (i.e., contralateral to right targets) to be less accurate and inconsistent/imprecise, and with higher lifts and more curved foot paths than stepping with the right leg (i.e., ipsilateral to right targets).

To further probe the nature of stepping actions using a one-item questionnaire of self-perceived difficulty/challenge, we conducted a frequency analysis on the survey question about the most challenging target per Choice. Based on the results of Experiment 1, we expected high self-perceived difficulty for lefts steps into posterior targets and to be scaled with target extent (i.e., far target the most difficult).

Design and data presentation and analysis were the same as in Experiment 1.

## Results

All twenty participants (10 men, mean ± SD, age = 27 ± 5 years, height = 1.72 ± 0.09 m, weight = 66.9 ± 12.3 kg) completed both Choice conditions without losing their balance and/or experiencing fatigue. After offline visual inspection of the videos, 8 (0.2%) and 27 (0.9%) trials out of 2880 were discarded from *Free* and *Constrained* conditions, respectively, because either the performer attempted to correct movement at the endpoint (see error correction arrow in **Fig. 4**), stepped before the ‘Go’ signal, or mistakenly used the dominant right limb in the *Constrained* condition. In contrast to Experiment 1, no participant used the contralateral leg to step into a target in *Free*. Findings from this experiment generally replicated the face validity, safety (i.e., participants completed all experimental procedures without any adverse events), and the methodological feasibility of the experimental paradigm and provided novel quantitative, kinematic evidence about stepping performance that pertain to the choice of stepping limb for goal-directed behaviors.

**Fig. 4.**
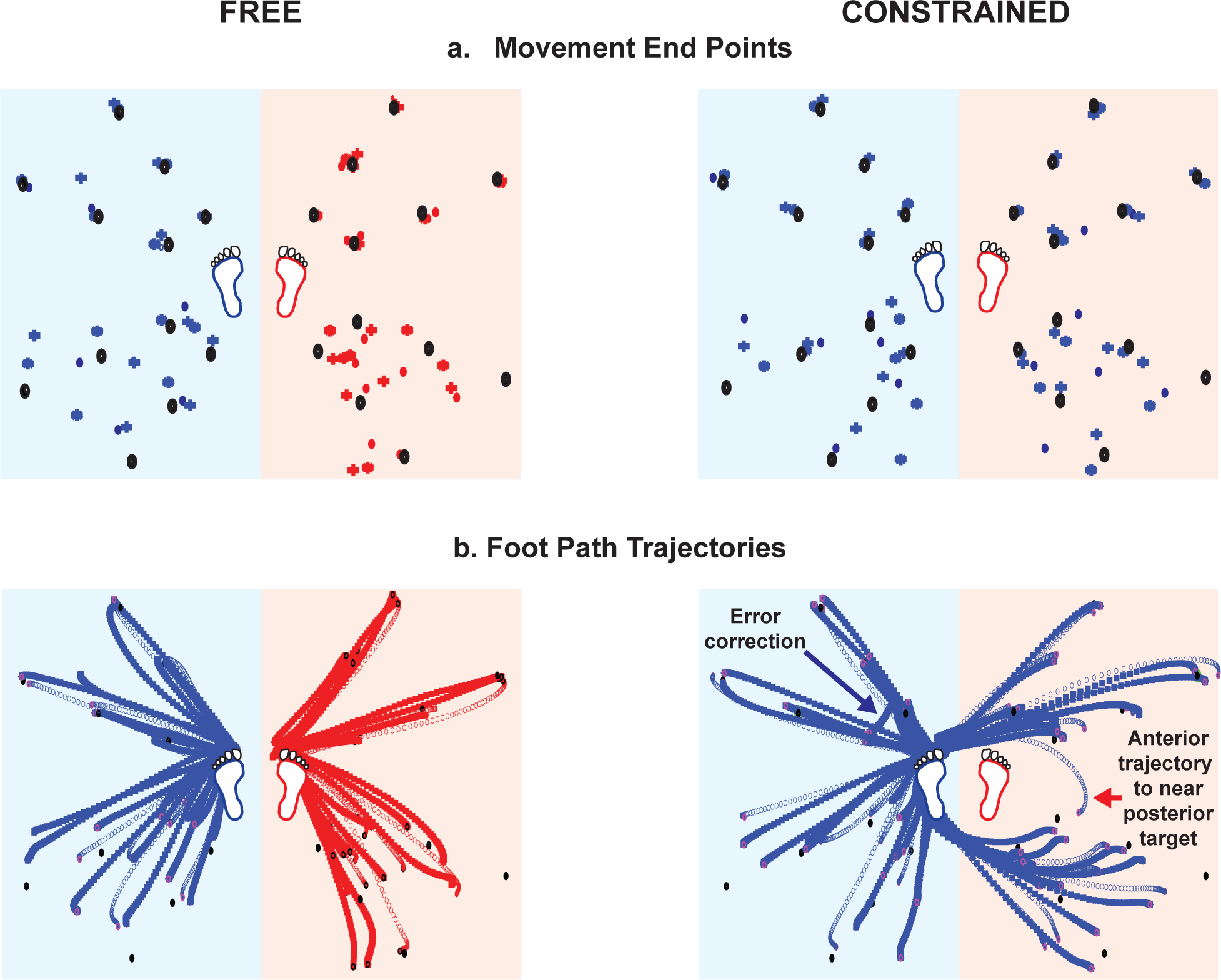
Representative trial data of movement end points (**a**) and foot path trajectories (**b**) relative to the targets during *Free* and *Constrained*. **(a)** Movement end points at each stepping trial during Free and Constrained; black filled ovals denote the targets, blue and red symbols (dot, cross, star and represent the three trials executed per target) denote the movement end points of the left and right stepping actions, respectively. **(b)** Foot path trajectories for each stepping trial during Free and Constrained. Blue and red lines denote the foot path trajectories of the left and right stepping actions, respectively. Black filled ovals denote the targets. Open pink and black circles indicate the movement end points during left and right stepping actions, respectively. One stepping trial during *Constrained* was an error correction (see blue arrow in the upper left quarter); it was removed from data analysis. Also, this participant stepped into one of the near posterior targets (i.e., green target in RP) by moving the left leg in front of the right leg for a single trial (see red arrow in the bottom right quarter), which was the first attempt out of three. **(a-b)** The footprints (blue: left leg; red: right leg) represent the start and end position before and after targeted stepping action. Blue and red shaded areas indicate the left and right hemispaces in which the left and right targets are located, respectively.

### Representative participant’s data

**Fig. 4** is a visual representation of trial data from one representative participant showing movement end points (**a**) and foot path trajectories (**b**) relative to each target during *Free* and *Constrained* conditions. The closer the movement end points were to the target (accuracy) and to each other (consistency/precision), the higher the success was. Similarly, the straighter the foot path trajectories (i.e., step path ratio closer to 1), the lower the motor effort was (note that foot lifts are not depicted).

#### Stepping actions into target regions using the ipsilateral limb

Movement end points (**Fig. 4a** – see the left steps/blue symbols in the light blue shaded area in *Free* and *Constrained* and right steps/red symbols in the light red shaded area in *Free*), for anterior stepping actions (i.e., forward stepping and presence of visual input) are relatively close to the target (i.e., high accuracy with slight overshoot) and with high consistency/precision across the three trials. Conversely, movement end points for posterior steps (i.e., backward stepping and absence of visual input) are relatively far from the target (i.e., low accuracy and undershot) and considerably inconsistent/imprecise across the three trials. Note that for anterior stepping, accuracy and consistency of the movement end points were similar across the three target extents, whereas for the posterior stepping both accuracy and consistency of the movement end points scaled with target extent (i.e., the farther the target extent the higher was the constant error and variable error). Independent of stepping direction (forward vs. backward), foot path trajectories to ipsilateral targets are relatively straight (**Fig. 4b** – see the left steps/blue paths in light blue shaded areas in *Free* and *Constrained* and right steps/red paths in light red shaded areas in *Free*).

#### Stepping actions into the right targets using the contralateral left limb

The pattern of the movement end points (**Fig. 4a** – see the left steps/blue symbols in the light red shaded right hemispace in *Constrained)* is like that using the ipsilateral right leg (**Fig. 4a** – see right steps/red symbols in the light red shaded hemispace in *Free*). Accuracy and consistency for anterior targets are higher than for posterior targets. Further, it appears that for anterior stepping, accuracy and consistency of the movement end points were similar across the three target extents, whereas for the posterior stepping both accuracy and consistency/precision decreased as target extent increased (i.e., higher constant and variable error). Like movement end points, foot path trajectories were dependent on stepping direction (**Fig. 4b –** see left steps/blue foot paths in the light red shaded right hemispace in *Constrained*). For anterior stepping, foot path trajectories were relatively straight, yet foot path trajectories were straighter as target distance increased. For posterior stepping, the foot path trajectories were markedly curved, yet note that foot path trajectories got straighter as target distance increased. For both anterior and posterior near targets, foot path trajectories were the most curved in large part because the contralateral stepping limb had to move around the right standing limb (**Fig. 4b –** see red footprint in the light red shaded right hemispace in *Constrained*).

### Group Data

#### Stepping Limb Use per Choice Condition

In *Free*, the limb choice results replicated those from Experiment 1 (not shown). Hence, the stepping limb was 100% of the time ipsilateral to the target. In *Constrained*, as designed and as in Experiment 1, all participants used their left leg to step onto all target regions and extents (not shown). Only for a few trials, four participants mistakenly used the right leg; these trials were removed from data analysis (see above). In all *Constrained* trials, participants stepped into the anterior and posterior right targets by moving the left leg in front of and behind the right standing leg, respectively. The only exception was the participant whose data are presented in **Fig.4** (see red arrow in *Constrained* in **Fig. 4b** right panel).

**Table 2** summarizes the statistical results for each kinematic measure for stepping actions into the left and right target regions in both Choice conditions; results of any post-hoc analyses are presented in the text.

**Table 2:** Summary of the results for the repeated measures ANOVA in Experiment 2.

| DVs/IVs | Choice<br>×<br>Target Region<br>×<br>Target Extent | Choice<br>×<br>Target Region | Choice<br>×<br>Target<br>Extent | Choice |
| --- | --- | --- | --- | --- |
| *Stepping into the LEFT Targets in Free and Constrained* |  |  |  |  |
| Constant Error (mm) | **$p=0.012$;**<br>**$F_{(1,2)}=4.99$** | $p=0.689$;<br>$F_{(1,2)}=0.17$ | **$p=0.023$;**<br>**$F_{(1,2)}=4.18$** | $p=0.373$;<br>$F_{(1,1)}=0.83$ |
| Variable Error (mm) | $p=0.211$;<br>$F_{(1,2)}=1.62$ | $p=0.520$;<br>$F_{(1,2)}=0.43$ | $p=0.937$;<br>$F_{(1,2)}=0.07$ | **$p=0.001$;**<br>**$F_{(1,2)}=14.29$** |
| Peak Vertical Foot Lift (mm) | $p=0.156$;<br>$F_{(1,2)}=1.95$ | $p=0.219$;<br>$F_{(1,2)}=1.62$ | $p=0.671$;<br>$F_{(1,2)}=0.40$ | $p=0.868$;<br>$F_{(1,2)}=0.023$ |
| Step Path Ratio | $p=0.740$;<br>$F_{(1,2)}=0.30$ | **$p=0.017$;**<br>**$F_{(1,2)}=6.85$** | $p=0.300$;<br>$F_{(1,2)}=1.25$ | **$p=0.002$;**<br>**$F_{(1,2)}=13.27$** |
| *Stepping into the RIGHT Targets in Free and Constrained* |  |  |  |  |
| Constant Error (mm) | **$p=0.004$;**<br>**$F_{(1,6)}=6.54$** | $p=0.914$;<br>$F_{(13)}=0.01$ | $p=0.075$;<br>$F_{(1,2)}=2.70$ | $p=0.697$;<br>$F_{(1,1)}=0.16$ |
| Variable Error (mm) | $p=0.693$;<br>$F_{(1,2)}=0.37$ | $p=0.585$;<br>$F_{(1,2)}=0.31$ | $p=0.657$;<br>$F_{(1,2)}=0.43$ | **$p=0.030$;**<br>**$F_{(1,2)}=5.49$** |
| Peak Vertical Foot Lift (mm) | **$p=0.0001$;**<br>**$F_{(1,2)}=20.06$** | **$p=0.012$;**<br>**$F_{(1,2)}=7.62$** | **$p=0.001$;**<br>**$F_{(1,2)}=43.59$** | **$p=0.001$;**<br>**$F_{(1,2)}=59.97$** |
| Step Path Ratio | $p=0.212$;<br>$F_{(1,2)}=1.62$ | **$p=0.001$;**<br>**$F_{(1,2)}=43.33$** | **$p=0.008$;**<br>**$F_{(1,2)}=5.60$** | **$p=0.001$;**<br>**$F_{(1,2)}=40.55$** |

#### Stepping into Left Target Regions (Table 2 top half; Fig 5-8a – left panels)

Given that the left leg (i.e., ipsilateral) was used in both Choice conditions, we expected neither interaction nor effect of Choice for any variable. Our expectations were supported only by one measure of effort. There was a significant 3-way interaction (Choice × Target Region × Target Extent) for constant error (unexpected), effect of Choice for variable error (unexpected), neither interaction nor effect of Choice for peak vertical foot lift (*p* > 0.05) (expected), and 2-way interaction (Choice × Target Region) for step path ratio (unexpected) (**Table 2** top half). In *Constrained* compared to *Free*, participants demonstrated greater accuracy (i.e. lower constant error; (t _(19)_ = 4.456; *p* = 0.0001; see purple asterisk in **Fig. 5a**) to the posterior near targets (**Fig. 1b II** – see the green targets in LP region), overall greater consistency/precision (i.e. lower variable error) (**Fig. 6a**), indifferent peak vertical foot lift (**Fig. 7a**), and slightly more curved trajectories (i.e. greater step path ratio) to anterior (LA) targets (t _(18)_ = –3.314; *p* = 0.004) (**Fig. 8a**). As expected, in *Free*, 17 of 20 participants perceived the most challenging target to be in the left posterior (LP) target region with the farthest extent coming in with the highest frequency (Near: 2, Middle: 4, Far: 11), whereas no participant perceived a left target to be the most challenging target in *Constrained*.

**Fig. 5.**
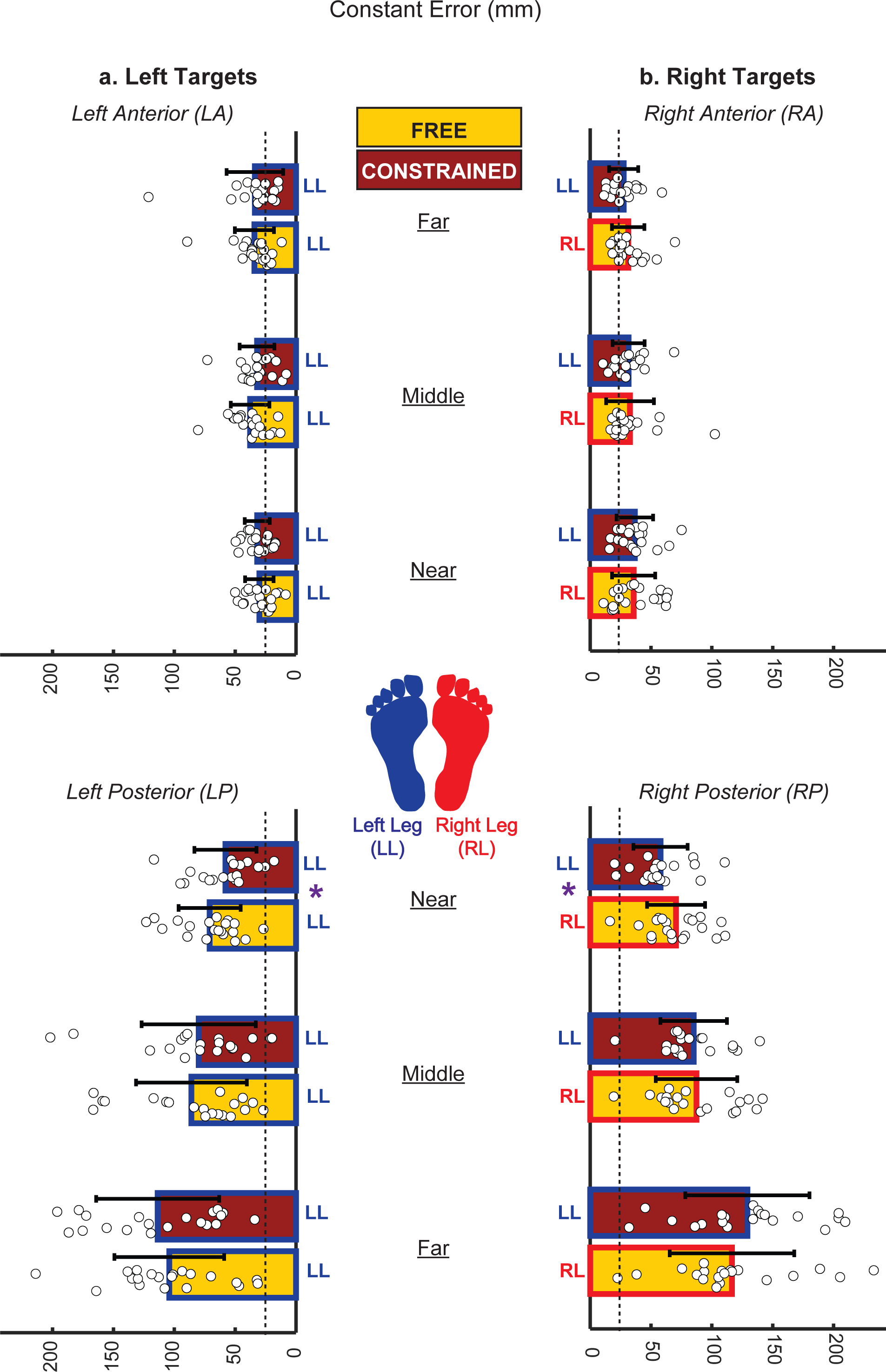
Constant error (mm) for stepping actions into the left target regions **(a)** and right target regions **(b)**. **(a & b)** White filled dots denote individual data points (N = 20) while the colored (Gold: *Free*; Cardinal: *Constrained*) bars and black error bars indicate group means ± 1 SD, respectively. The vertical dashed lines indicate the radius of the target (i.e., 25 mm); therefore, any dots lower than those dashed lines denote that these participants stepped within the target (i.e., goal of the task). Independent of stepping limb, note that in both anterior targets (LA and RA), many participants stepped accurately into the target (i.e., most dots lower than the dashed line) whereas for the posterior target regions (LP and RP) only a few participants stepped accurately into the target (i.e., only a few dots lower than the dashed line). Blue and red outlined bars denote left (LL) and right leg (RL) were used, respectively. Purple asterisk (*) denotes significance in post-hoc test (corrected p-value) only if a significant (p<0.05) Choice × Target Region × Target Extent interaction was found.

**Fig. 6.**
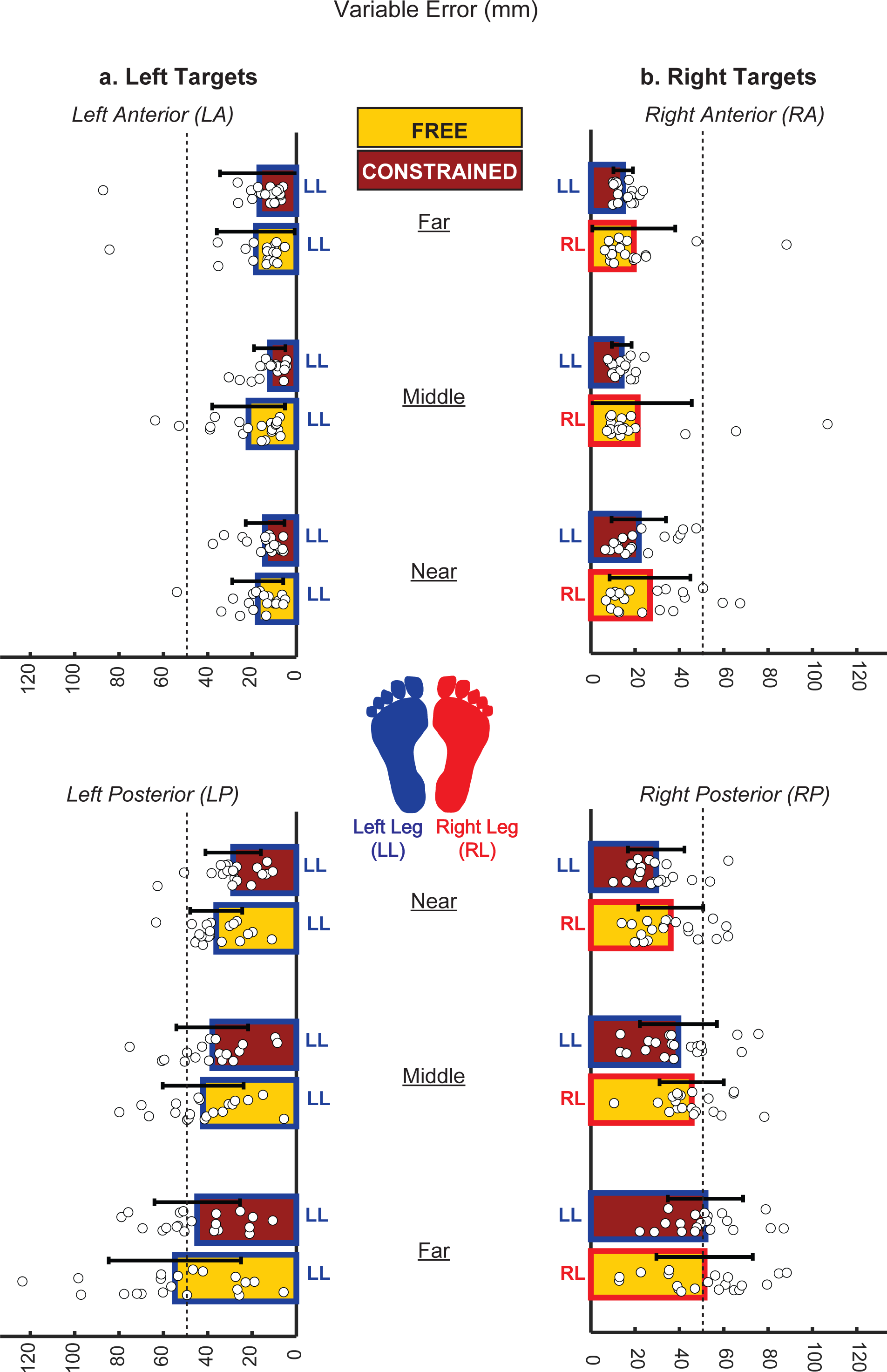
Variable error (mm) for stepping actions into the left target regions **(a)** and right target regions **(b)**. **(a & b)** White filled dots denote individual data points (N = 20) while the colored (Gold: *Free*; Cardinal: *Constrained*) bars and black error bars indicate group means ± 1 SD, respectively. Blue and red outlined bars denote left (LL) and right leg (RL) were used, respectively. The vertical dashed lines indicate the diameter of the target (i.e., 50 mm). Any dots lower than those dashed lines denote that these participants stepped with high consistency within target diameter. Note that in both anterior target regions (LA and RA), majority of the participants stepped with high consistency (i.e., most dots lower than the dashed line), especially for stepping actions in *Constrained* (i.e., all dots are lower than the dashed line and scaled with target extent with an exception one participant in LA target region and far target extent). For the posterior target regions (LP and RP) nearly half of the participants stepped with high consistency (i.e., most dots lower than the dashed line; Constrained > Free) which scaled with target extent (i.e., consistency decreased from near to far target extent; near > middle > far).

**Fig. 7.**
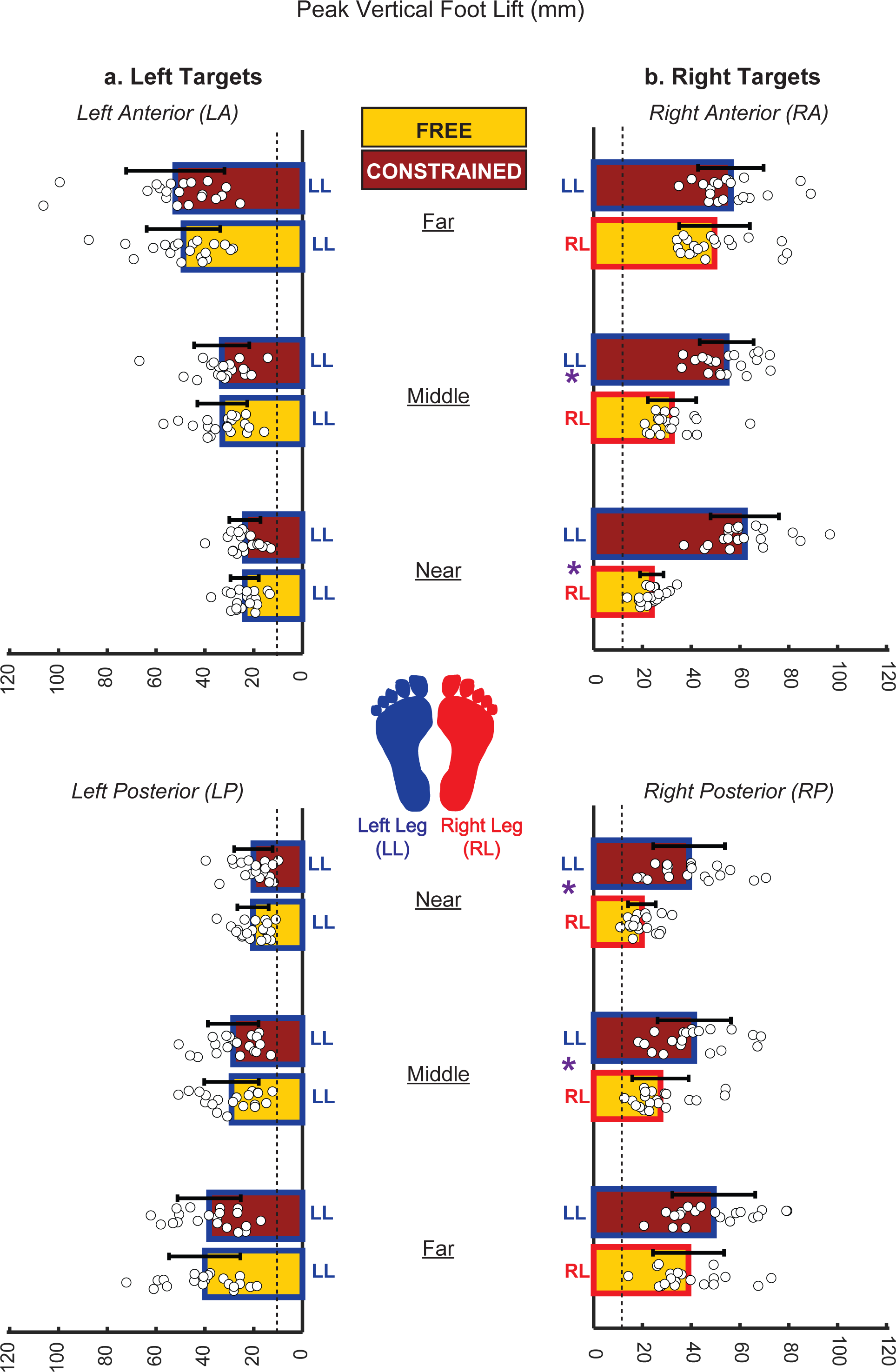
Peak vertical foot lift (mm) for stepping actions into the left target regions **(a)** and right target regions **(b)**. **(a & b)** White filled dots denote individual data points (N = 20) while the colored (Gold: *Free*; Cardinal: *Constrained*) bars and black error bars indicate group means ± 1 SD, respectively. Blue and red outlined bars denote left (LL) and right leg (RL) were used, respectively. The vertical dashed lines indicate the invariant toe clearance (12.7 mm) during mid-swing phase of walking reported in young adults (Winter et al. 1990). Any dots close (± 6 mm) to this dashed line (12.7 mm) implies that participants adapt similar kinematic pattern as during swing, whose pattern resembles the stepping action in this study. For all ipsilateral steps (LL in left targets and RL in right target regions), note that the peak vertical foot lifts increased with target extent, whereas for the contralateral steps (LL in right target regions), note that the peak vertical foot lifts were independent of target extent. Purple asterisk (*) denotes significance in post-hoc test (corrected p-value) only if a significant (p<0.05) Choice × Target Region × Target Extent interaction was found.

**Fig. 8.**
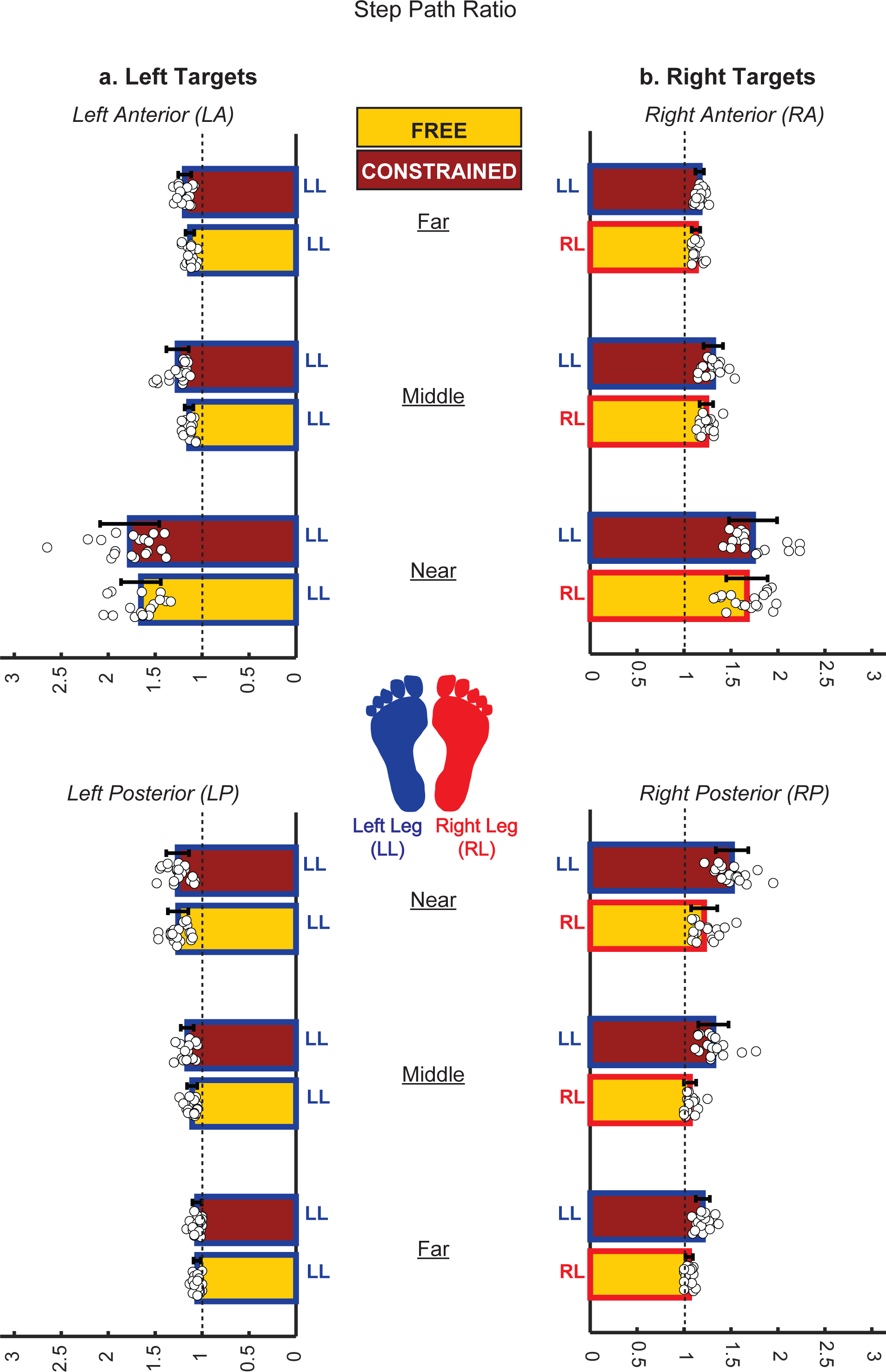
Step path ratio for stepping actions into the left target regions **(a)** and right target regions **(b)**. **(a & b)** White filled dots denote individual data points (N = 20) while the colored (Gold: *Free*; Cardinal: *Constrained*) bars and black error bars indicate group means ± 1 SD, respectively. Blue and red outlined bars denote left (LL) and right leg (RL) were used, respectively. The vertical dashed lines indicate straight path (i.e., a ratio of 1: foot path distance equals movement distance); note that independent of stepping limb (i.e., Choice condition) and target regions, participants stepped relatively with straight path as the target extent increased. Also, note that participants stepped with relatively curved paths to the near targets, especially those located anteriorly (independent of Choice condition).

#### Stepping into Right Target Regions (Table 2 bottom half; Fig 5-8b – right panels)

Given that a different leg was used for each Choice condition, we expected either an interaction or effect of Choice condition for all variables. Our expectations were supported by both measures of success and effort. Hence, there was a significant 3-way interaction (Limb Choice × Target Region × Target Extent) for constant error, effect of Choice for variable error, 3-way interaction (Choice × Target Region × Target Extent) for peak vertical foot lift, and 2-way Choice × Target Region and Limb Choice × Target Extent interactions on step path ratio (**Table 2** bottom half). There was no significant difference in constant error between the two Choice conditions for the right anterior targets. However, as seen for stepping actions into left targets using the ipsilateral left leg, in the *Constrained* compared to *Free*, participants demonstrated greater accuracy (i.e., lower constant error; t _(19)_ = 2.884; *p* = 0.010; see purple asterisk in **Fig. 5b**) only for the posterior near targets (**Fig. 1b III** – see the green targets in RP region). For variable error, in *Constrained* compared to *Free*, participants demonstrated overall greater consistency/precision (i.e., lower variable error; see *Constrained* bars that in general are shorter than *Free* bars in **Fig. 6b**). For peak vertical foot lift, in *Constrained* compared to *Free* trials for all right targets, participants demonstrated higher vertical foot lift for near and middle targets in both anterior (Near: t _(19)_ = –12.065; *p* = 0.0001; Middle: t _(19)_ = –8.534; *p* = 0.0001) and posterior (Near: t _(19)_ = –6.468; *p* = 0.0001; Middle: t _(19)_ = –3.917; *p* = 0.001) target regions (see the purple asterisks in **Fig. 7b)**. For step path ratio, in *Constrained* compared to *Free* trials for all right targets, participants demonstrated more curved step paths (i.e., higher step path ratio) for the posterior (RP) target region (t _(18)_ = – 8.743; *p* = 0.0001) and for all target extents (Near: t _(18)_ = –4.910; *p* = 0.0001; Middle: t _(18)_ = – 5.893; *p* = 0.0001; Far: t _(18)_ = –6.438; *p* = 0.0001) **(Fig. 8b**). Finally, in *Free,* only 3 of 20 participants perceived the most challenging target to be in the right posterior (RP) region and at the farthest extent. Not surprisingly, all 20 participants after *Constrained* trials, identified the most challenging target to be also in the right posterior (RP) region with the farthest extent coming in with the highest frequency (i.e., Near: 2, Middle: 3; Far: 15).

## Discussion

Through two early-stage phased experiments, we explored whether expected success and/or motor effort play a role in the motor decision-making of limb choice during a volitional goal-directed stepping in neurotypical adults. Given the experimental design, which allowed leg choice manipulation, the ipsilateral left leg was used in both Choice conditions (*Free* and *Constrained*) to step into left targets. Therefore, stepping actions into the left targets were used as a proxy for neurotypical limb selection without extrinsic constraints; we expected participants would step into the ipsilateral left targets with high success (i.e., low constant and variable error) and relatively low motor effort (low vertical foot lifts and straight paths) in both Choice conditions. Conversely, left steps into the contralateral right targets were used as an artificial proxy for intrinsic constraint, such as hemiparesis where the task is more difficult and postural stability is challenged. Therefore, when the Choice condition was the *Constrained* condition, we expected that success and effort would decrease and increase, respectively.

Our expectations were partially supported but with a few unexpected findings. Overall, there were four novel and robust findings. First (*expected finding*), when free to choose, participants uniformly selected the limb ipsilateral to the target (i.e., hemispace bias) and with slight leg dominance bias for central targets. Second (*unexpected finding*), success measures (i.e., constant error and variable error) provide evidence that accuracy and consistency/precision for stepping actions can improve with a short bout of random practice (Shea and Morgan 1979) to both anterior (variable error in all target extents) and posterior (constant error in near targets and variable error in all target extents) target regions independent of Choice condition. Third (*expected finding*), motor effort measures (i.e., vertical peak foot lift and step path ratio) provide evidence that effort for stepping actions into left and right targets may be an important influence on decision-making about which limb to choose for stepping; hence, motor effort is greater when the leg contralateral to the target is used (i.e., vertical peak foot lift for the near and middle right anterior and posterior targets; step path ratio for posterior targets and across target extents). Fourth (*unexpected finding*), participants perceived those posterior targets (mainly the farthest away) to be the most difficult primarily during non-dominant left steps independent of Choice condition. In the following sections we unpack, discuss, and provide plausible explanations both for the expected and unexpected findings.

### Limb selection revealed hemispace bias for ipsilateral targets

In *Free*, the robust finding that the leg ipsilateral to the target was chosen 100% of the time not only confirms the hemispace bias effect for discrete stepping, but it replicates the arm choice behavior studies in both neurotypical adults and neurologically impaired individuals (Helbig and Gabbard 2004; Coelho et al. 2013; Han et al. 2013).

Taken together, goal-directed actions may assume similar motor control strategies independent of effector (arm vs. leg) (Brunt et al. 2000; Sparrow et al. 2003). Georgopoulos and Grillner (1989) have postulated that despite task-related differences reaching and locomotion may share similar neural mechanisms across the neuromotor axis. They discussed that some species of non-human primates, who are capable of both quadrupedal and bipedal walking, can use their forelimbs for reaching and grasping a tree branch to navigate through the forest. Specifically, this action demands high precision in positioning the hand and timely accurate, fast, and coordinated alternative grasping using the right and left hand. As in non-human primates, reaching and walking in cats also share similar motor cortical and subcortical mechanisms (Dyson et al. 2014; Yakovenko and Drew 2015). Furthermore, to meet those action demands, sensory information is crucial, particularly vision (Georgopoulos 1986; Drew 1991; Drew et al. 2008). During goal-directed actions, the motor system uses vision for visual localization of the object in space to which the limb must move, visual monitoring of the limb right before and during movement, and visual adjustment of the limb at the final position (Georgopoulos 1986; Scott et al. 2015). Given the similarities in visuomotor coordination inherent to both tasks, stepping and reaching likely share similar mechanisms of limb selection.

During reaching, participants used the ipsilateral arm to the target in most trials, but not in all (i.e., the contralateral arm was used in some trials as well) (Helbig and Gabbard 2004; Coelho et al. 2013). Additionally, for reaching contralateral targets, the dominant arm is used more frequently than the non-dominant arm. The findings for those studies contrast with the present study, in which all participants always utilized their ipsilateral leg for the left and right targets whereas leg dominance had no role in stepping to ipsilateral targets. Therefore, the robust hemispace bias demonstrated in the present study suggests that it might be stronger for stepping than for reaching. Yet, the remaining question is why so?

One explanation could be that for unimanual arm tasks the moving arm can operate relatively independently from the non-moving arm. This is not the case for stepping; the performance of the moving limb depends on the dynamic stability capacity of the standing limb (Lyon and Day 1997). Therefore, crossing the midline with the reaching arm is a frequent motor pattern (especially for the dominant arm) and feasibly easy, whereas crossing the midline with the stepping limb is not a frequent pattern and entails non-zero postural challenges.

Another explanation for this strong hemispace bias is that leg dominance may not be as strong as arm dominance. It has been long debated how leg dominance can be characterized and whether it exists to the same degree as for the hand (Peters 1988). Therefore, taken together, it is likely that our participants demonstrated a relatively strong hemispace bias in order to ensure dynamic postural stability, take advantage of target proximity, and to secure a comfortable end-state position (Rosenbaum et al. 1990; Rosenbaum et al. 1996) while successfully executing the task with relatively low effort. Further investigation is warranted to tease apart the independent contribution of these factors that together contribute to such a strong hemispace bias for stepping actions.

### Success measures provide evidence that accuracy and precision for stepping actions can improve with a short bout of random practice to both anterior and posterior targets

Surprisingly, both accuracy and precision had the exact same pattern of results for left and right targets, independent of Choice condition. Specifically, for both ipsilateral left steps into left targets and contralateral left steps into right targets, stepping was more successful (i.e., both accuracy and consistency/precision increased) in the second block (i.e., *Constrained*) compared to the first block (i.e., *Free*) of randomly ordered trials.

Participants stepped into left posterior near targets (see green targets located in LP in **Fig. 1**) with higher accuracy (i.e., lower constant error) during *Constrained* than during *Free*. Note that these targets are right behind the stepping leg with no view of their location or stepping limb; visual information regarding target location and moving limb affect accuracy and precision during a stepping task (Rietdyk and Drifmeyer 2010; Smid and den Otter 2013). Given the extent and location of these targets, participants may have improved their accuracy just because this extent might be the most employed during protective stepping responses and could adapt after only three trials (Experiment 2). Of note, in *Constrained* trials, participants were more accurate stepping into posterior near targets without changing either foot lift or foot path trajectory (see the motor effort data of both *Choice* conditions for the near targets in the left bottom panels – LP in **Fig 7a** and **Fig 8a**). In addition, and in contrast to our expectations, precision was higher (i.e., lower variable error) during *Constrained* than during *Free* (i.e., independent of target region and extent). As with accuracy, precision may increase after only three randomly assigned trials of these more familiar stepping tasks, reflecting refined sensorimotor adaptation in neurotypical adults (Bastian 2008). Taken together, neurotypical adults can step into ipsilateral targets located in various regions and extents with relative success, which increases with random order practice (Shea and Morgan 1979).

For the contralateral left and ipsilateral right steps into right targets, the *exact same* counter-intuitive results for constant error and variable error were observed as for ipsilateral stepping into left targets; greater success in *Constrained* compared with *Free*. Hence, left stepping actions were more accurate (i.e., lower constant error) for the right posterior near targets (see green targets located in RP in **Fig. 1**) and more precise (lower variable error) across all right targets compared to right steps into the ipsilateral right targets. As previously, one possible explanation might simply be that neurotypical young adults can increase their stepping precision and accuracy within 6 trials that are randomly ordered (24 targets x 3 trials per *Choice* condition) and with the more difficult of the two conditions (i.e. Constrained) following the easier condition (i.e. Free) regardless of whether the stepping limb is either ipsilateral or contralateral. This simple explanation cannot be ruled out. Of note, the more difficult and challenging *Constrained* condition may elicit more attention and more motivation thereby provoking greater goal-action coupling (Wulf and Lewthwaite 2016). The target specificity (i.e., posterior near targets) for the accuracy results are consistent with that observed for the ipsilateral left stepping actions. It is also important to point out that participants were accurate and precise during these contralateral steps by lifting the foot higher and moving the foot with more curved paths (see the motor effort data of both *Choice* conditions in the right bottom panels – RP in **Fig 7b** and **Fig 8b**). Therefore, neurotypical adults might be capable of maintaining successful stepping by manipulating the effort required to perform those stepping movements.

The difference between the precision/consistency and accuracy results – precision being generalized across all targets (left in Free trials ad right in Constrained trials) and accuracy confined to the posterior near targets (left in Free trials ad right in Constrained trials) – might reflect the greater sensitivity of the precision measure (i.e. variable error) compared with the accuracy measure (i.e. constant error). It is a well-known phenomenon in motor learning that variable error continues to decline with practice even when constant error has plateaued (Schmidt et al. 2019). Given that these stepping actions are relatively well-practiced behaviors, any benefit of a short bout of random practice could show up across the board using the more sensitive variable error measure, but only for the most challenging posterior targets using the less sensitive constant error measure.

### Effort measures support the notion that the expected effort for stepping actions into left and right targets may influence the choice of stepping limb

Participants moved their left leg into the ipsilateral left anterior (LA) targets with greater curvature in *Constrained* compared with *Free* (primarily for near targets – see top left panel – LA in **Fig. 8a**). Given that for the anterior target region participants demonstrated greater precision (i.e., lower variable error) during Constrained, the increase of curvature could be an adaptive strategy to ensure end point precision relative to the target. In addition, anterior targets can make use of vision to increase precision in this more challenging condition. However, the step path ratio was not different between conditions for the ipsilateral left posterior (LP) targets. One possible explanation is that the adaptive strategy described for anterior targets was one that could make use of visual input about target location. Importantly, this strategy was not possible for posterior targets where participants had to rely on remembered target location.

For contralateral left and ipsilateral right steps into right targets, not surprisingly, participants stepped into the right targets in general with greater motor effort using the contralateral left leg compared with the ipsilateral right leg. In particular, and compared with ipsilateral right leg steps, contralateral left leg steps exhibited a greater peak vertical foot lift for near and middle targets in both right target regions. Similarly, they stepped with more curved paths for the posterior targets and across target extents using the left leg than the right leg. On the surface, these findings are not surprising; therefore, we present four plausible explanations. First, the higher vertical lifts and curved stepping paths are likely necessitated by having to move around the right standing limb. Second, the relatively greater visual uncertainty (i.e., lower visual inputs of the targets) of these targets may necessitate greater effort and attention (Cohen et al. 2010). Third, the low motor familiarity (i.e., unnatural, and infrequent movement), especially crossing the stepping limb in front of the standing limb (unless we are considering, for example, an elite ballet dancer who performs such movements frequently). Finally, the high postural stability required of the right standing limb while the contralateral moving leg is translated outside of the base of support (Day and Bancroft 2018).

### Participants perceived posterior targets (mainly the farthest) to be the most difficult during non-dominant left steps

For ipsilateral left steps into left targets, 17 out of 20 participants in Experiment 2 identified targets located in the left posterior (LP) region to be the most difficult whereas no participant perceived left target region (neither LA nor LP) to be difficult during *Constrained* trials. Of those 17 who perceived the left posterior targets to the most difficult, eleven participants (65%) a target located in the farthest extent. For the ipsilateral right steps and the contralateral left steps to right targets in *Free* and *Constrained* trails, respectively, only three participants selected a right posterior target to be difficult in Free whereas all twenty participants (100%) selected a right posterior target to be the most difficult in Constrained trials. For both Choice conditions, those selected right posterior targets were mainly in the farthest extent (Constrained: 75%; Free: 100%).

Taken together, these findings suggest that the perceived difficulty of a target may depend on leg dominance (i.e., non-dominant), stepping direction (backward stepping), and target extent (farthest away). During voluntary stepping, non-dominant leg is usually selected the standing limb whereas the dominant leg is usually selected as the stepping limb (Peters 1988); therefore, using the non-dominant leg for stepping may increase the perceived difficulty of the target regardless of its location (ipsilaterally or contralaterally). Further, in contrast to forward stepping, postural stability is challenged because of limited vision and the biomechanical challenges of backward voluntary stepping (Winter et al. 1989; Lee et al. 2014). Also, increase of the moving distance (target extent) required reaches further beyond the base of support (i.e., greater variability of center of pressure) and longer single standing limb durations (Duarte and Freitas 2005; Duarte and Latash 2007; Oku et al. 2023); therefore, target extent may also influence how neurotypical adults perceive target difficulty. These findings strengthen the notion that stepping backwards to far targets with limited to no online visual input and with the non-dominant limb is perceived to be difficult by neurotypical young adults.

### Success vs. Effort: Which has greater influence on the choice of the stepping limb?

The present findings from the success, effort, and perceived difficulty measures provide initial evidence for the roles played by success and/or effort in stepping limb selection. Taken together, the four novel findings suggest that neurotypical adults may select the stepping limb based primarily on effort and to a lesser extent based on success. This is inconsistent with findings from studies of limb selection for upper extremity aiming in which both factors were found to influence arm choice in neurotypical adults and stroke survivors (Schweighofer et al. 2015; Hirayama et al. 2022; Kim et al. 2022; Nguyen et al. 2023). This discrepancy between aiming and stepping might be explained by fundamental differences in task demands alluded to earlier and discussed further below. Contrary to arm reaching, postural stability is a vital component of stepping behavior. The standing limb must be capable of providing a stable base of support (i.e. dynamic balance) while the stepping limb is used to step to a target accurately(Duarte and Freitas 2005; Duarte and Latash 2007; Day and Bancroft 2018). It makes sense that for stepping tasks, minimizing effort may enable the necessary postural stability for task success. Yet, no participant lost balance for the left steps into the contralateral right targets; so postural stability was maintained by the right standing limb, but at the cost of motor effort (expended by the stepping limb). Though we did not quantify the effort associated with postural stability here, we argue that postural stability for steps into targets with higher challenge (i.e., lefts steps to contralateral targets) would be challenged (e.g., higher displacement of the center of pressure) (Duarte and Freitas 2005; Duarte and Latash 2007).

Further mention is warranted of fundamental differences in task requirements and what constitutes success for aiming and stepping tasks. We speculate that successful stepping requires accuracy, but also importantly, postural stability to avoid loss of balance during the step—maintenance of postural stability requires motor effort. By contrast, successful aiming depends to a greater degree on the accuracy of the endpoint (especially for grasping an object) than on motor effort as defined for postural control. The motor effort for aiming movements has been typically associated with the generation of fast movements (Schweighofer et al. 2015; Kim et al. 2022). Further research is needed to unravel these factors and with a head-to-head comparison including additional measures of postural stability that are beyond the scope of this preliminary investigation.

### Methodological Considerations

Four methodological considerations must be noted when interpreting our preliminary findings from these two early-stage experiments. First, both experiments had an asymmetric design in terms of the leg used for *Constrained* trials; by design it was always the left non-dominant leg. Therefore, we cannot definitively conclude whether leg dominance influences the choice of stepping limb. Accordingly, such an investigation (i.e., symmetric design of the leg used for *Constrained* trials) would be relevant particularly for studies of clinical cohorts for which the dominant limb can be impaired. Second, we did not counterbalance the two Choice conditions. Rather we reasoned that having the *Free* condition first would be best for obtaining an unbiased evaluation of choice before attempting the *Constrained* condition where there was no choice. This blocked design, with the more difficult condition following the easier one, may have been the optimal schedule along with a within-block random order target arrangement for eliciting improved performance over practice. We did indeed see the benefits of a random practice schedule for both success measures in Experiment 2 (i.e., *Constrained* trials were more accurate and consistent/precise than *Free* trials for both ipsilateral and contralateral left steps). Third, we calculated bilateral position-derived kinematics, but we did not acquire each supporting foot’s kinetics (e.g., center of pressure), which could have been used as a proxy of postural stability (Winter 1995). Specifically, we could have quantified the postural demand and tested the hypothesis that the greater the center of pressure displacement or velocity, the greater the postural instability (Winter 1995). Using such a kinetic measure would be useful for providing quantitative evidence on the postural challenges/demands of stepping, especially for volitional stepping into targets located in awkward locations (e.g., contralateral) and extents (i.e., far) relative to the stepping limb and with limited vision (Inaba et al. 2020). Fourth, our design pertains to the fact that target extent was fixed and not normalized to each participant’s leg length; however, we did select a range of target extents that we thought would accommodate a fairly large range of different leg lengths.

## Conclusion

In this set of experiments, we explored whether success and/or motor effort have a role in the decision-making for stepping limb choice during a volitional goal-directed stepping task. We conclude that neurotypical adults demonstrated a robust hemispace bias in stepping actions as well as slight leg dominance bias for the central anteroposterior targets and greater influence of motor effort on the choice of stepping limb than actual success. The present findings provide an initial foundation for future work that should probe these findings’ robustness in neurological cohort with unilateral paresis (intrinsic constraint) to advance our understanding of motor decision processes for goal-directed mobility behaviors.

## Acknowledgements

The authors would like to thank Dr. Nicolas Schweighofer, PhD for his useful inputs during the development of the task, Dr. Rini Varghese, PT, PhD for insightful comments during early manuscript preparation, all participants, and the staff of CATZ training center, Pasadena, CA, for their assistance with data collection.

## Author Contributions

The resources for the data acquisition and analysis were provided by CJW, who supervised the study. The conception and the experimental design of the study were developed by CCC, YHL, and CJW. Experiments were performed by CCC and YHL. Data were acquired by CCC, YHL, EREW, GMC, and MG. Codes for data analyses were developed by CCC and EREW. Data were curated by CCC, YHL, MG and analyzed by CCC. Data were visualized and results were validated by CCC and CJW. Article was drafted by CCC and edited and revised by CCC and CJW. The final version of the manuscript was read and approved by CCC, EREW, GMC, MG, YHL, and CJW.

## Data Availability

The data that support the findings from this study are available from the corresponding author, upon reasonable request.

## References

1. Bastian AJ (2008) Understanding sensorimotor adaptation and learning for rehabilitation. Curr Opin Neurol 21:628–633 doi: 10.1097/WCO.0b013e328315a293

2. Berchicci M, Russo Y, Bianco V, et al. (2020) Stepping forward, stepping backward: a movement-related cortical potential study unveils distinctive brain activities. Behav Brain Res 388:112663 doi: 10.1016/j.bbr.2020.112663

3. Brunt D, Short M, Trimble M, Liu SM (2000) Control strategies for initiation of human gait are influenced by accuracy constraints. Neurosci Lett 285:228–230

4. Bryden PJ, Roy EA (2006) Preferential reaching across regions of hemispace in adults and children. Dev Psychobiol 48:121–132 doi: 10.1002/dev.20120

5. Calvert GA, Bishop DV (1998) Quantifying hand preference using a behavioural continuum. Laterality 3:255–268 doi: 10.1080/713754307

6. Cisek P (2012) Making decisions through a distributed consensus. Curr Opin Neurobiol 22:927–936 doi: 10.1016/j.conb.2012.05.007

7. Coelho CJ, Przybyla A, Yadav V, Sainburg RL (2013) Hemispheric differences in the control of limb dynamics: a link between arm performance asymmetries and arm selection patterns. J Neurophysiol 109:825–838 doi: 10.1152/jn.00885.2012

8. Cohen RG, Biddle JC, Rosenbaum DA (2010) Manual obstacle avoidance takes into account visual uncertainty, motor noise, and biomechanical costs. Exp Brain Res 201:587–592 doi: 10.1007/s00221-009-2042-8

9. Cortes JC, Goldsmith J, Harran MD, et al. (2017) A Short and Distinct Time Window for Recovery of Arm Motor Control Early After Stroke Revealed With a Global Measure of Trajectory Kinematics. Neurorehabil Neural Repair 31:552–560 doi: 10.1177/1545968317697034

10. Cos I, Belanger N, Cisek P (2011) The influence of predicted arm biomechanics on decision making. J Neurophysiol 105:3022–3033 doi: 10.1152/jn.00975.2010

11. Day BL, Bancroft MJ (2018) Voluntary steps and gait initiation. Handb Clin Neurol 159:107–118 doi: 10.1016/b978-0-444-63916-5.00006-9

12. Dessery Y, Barbier F, Gillet C, Corbeil P (2011) Does lower limb preference influence gait initiation? Gait Posture 33:550–555 doi: S0966-6362(11)00022-1 [pii] 10.1016/j.gaitpost.2011.01.008

13. Dominguez-Zamora FJ, Marigold DS (2021) Motives driving gaze and walking decisions. Curr Biol 31:1632–1642 e1634 doi: 10.1016/j.cub.2021.01.069

14. Drew T (1991) Visuomotor coordination in locomotion. Curr Opin Neurobiol 1:652–657

15. Drew T, Andujar JE, Lajoie K, Yakovenko S (2008) Cortical mechanisms involved in visuomotor coordination during precision walking. Brain Res Rev 57:199–211 doi: 10.1016/j.brainresrev.2007.07.017

16. Duarte M, Freitas SM (2005) Speed-accuracy trade-off in voluntary postural movements. Motor Control 9:180–196 doi: 10.1123/mcj.9.2.180

17. Duarte M, Latash ML (2007) Effects of postural task requirements on the speed-accuracy trade-off. Exp Brain Res 180:457–467 doi: 10.1007/s00221-007-0871-x

18. Duncan PW, Weiner DK, Chandler J, Studenski S (1990) Functional reach: a new clinical measure of balance. J Gerontol 45:M192–197

19. Dyson KS, Miron JP, Drew T (2014) Differential modulation of descending signals from the reticulospinal system during reaching and locomotion. J Neurophysiol 112:2505–2528 doi: 10.1152/jn.00188.2014

20. Faraji S, Wu AR, Ijspeert AJ (2018) A simple model of mechanical effects to estimate metabolic cost of human walking. Sci Rep 8:10998 doi: 10.1038/s41598-018-29429-z

21. Gallivan JP, Chapman CS, Wolpert DM, Flanagan JR (2018) Decision-making in sensorimotor control. Nat Rev Neurosci doi: 10.1038/s41583-018-0045-9

22. Georgopoulos AP (1986) On reaching. Annu Rev Neurosci 9:147–170 doi: 10.1146/annurev.ne.09.030186.001051

23. Georgopoulos AP, Grillner S (1989) Visuomotor coordination in reaching and locomotion. Science 245:1209–1210

24. Georgopoulos AP, Kalaska JF, Massey JT (1981) Spatial trajectories and reaction times of aimed movements: effects of practice, uncertainty, and change in target location. J Neurophysiol 46:725–743 doi: 10.1152/jn.1981.46.4.725

25. Gordon J, Ghilardi MF, Ghez C (1994) Accuracy of planar reaching movements. I. Independence of direction and extent variability. Exp Brain Res 99:97–111

26. Han CE, Kim S, Chen S, et al. (2013) Quantifying arm nonuse in individuals poststroke. Neurorehabil Neural Repair 27:439–447 doi: 10.1177/1545968312471904

27. Helbig CR, Gabbard C (2004) What determines limb selection for reaching? Res Q Exerc Sport 75:47–59 doi: 10.1080/02701367.2004.10609133

28. Hirayama K, Ito Y, Takahashi T, Osu R (2022) Relevant factors for arm choice in reaching movement: a scoping review. J Phys Ther Sci 34:804–812 doi: 10.1589/jpts.34.804

29. Hoffman M, Schrader J, Applegate T, Koceja D (1998) Unilateral postural control of the functionally dominant and nondominant extremities of healthy subjects. J Athl Train 33:319–322

30. Inaba Y, Suzuki T, Yoshioka S, Fukashiro S (2020) Directional Control Mechanisms in Multidirectional Step Initiating Tasks. Front Hum Neurosci 14:178 doi: 10.3389/fnhum.2020.00178

31. Keetch KM, Schmidt RA, Lee TD, Young DE (2005) Especial skills: their emergence with massive amounts of practice. J Exp Psychol Hum Percept Perform 31:970–978 doi: 10.1037/0096-1523.31.5.970

32. Kim HE, Avraham G, Ivry RB (2021) The Psychology of Reaching: Action Selection, Movement Implementation, and Sensorimotor Learning. Annu Rev Psychol 72:61–95 doi: 10.1146/annurev-psych-010419-051053

33. Kim S, Han CE, Kim B, Winstein CJ, Schweighofer N (2022) Effort, success, and side of lesion determine arm choice in individuals with chronic stroke. J Neurophysiol 127:255–266 doi: 10.1152/jn.00532.2020

34. Kim S, Park H, Han CE, Winstein CJ, Schweighofer N (2018) Measuring Habitual Arm Use Post-stroke With a Bilateral Time-Constrained Reaching Task. Front Neurol 9:883 doi: 10.3389/fneur.2018.00883

35. Lee PY, Gadareh K, Bronstein AM (2014) Forward-backward postural protective stepping responses in young and elderly adults. Hum Mov Sci doi: 10.1016/j.humov.2013.12.010

36. Lyon IN, Day BL (1997) Control of frontal plane body motion in human stepping. Exp Brain Res 115:345–356 doi: 10.1007/pl00005703

37. Mamolo CM, Roy EA, Rohr LE, Bryden PJ (2006) Reaching patterns across working space: the effects of handedness, task demands, and comfort levels. Laterality 11:465–492 doi: 10.1080/13576500600775692

38. McLlory WE, Maki BE (1996) Influence of Destabilization on the Temporal Characteristics of “Volitional” Stepping. J Mot Behav 28:28–34 doi: 10.1080/00222895.1996.9941730

39. Nguyen H, Phan T, Shadmehr R, Lee SW (2023) Choice of Arm Use in Stroke Survivors is Largely Driven by the Energetic Cost of the Movement. Neurorehabil Neural Repair 37:183–193 doi: 10.1177/15459683231164788

40. Oku K, Tanaka S, Kida N (2023) Direction and distance dependency of reaching movements of lower limb. PLoS One 18:e0290745 doi: 10.1371/journal.pone.0290745

41. Peters M (1988) Footedness: asymmetries in foot preference and skill and neuropsychological assessment of foot movement. Psychol Bull 103:179–192 doi: 10.1037/0033-2909.103.2.179

42. Przybyla A, Coelho CJ, Akpinar S, Kirazci S, Sainburg RL (2013) Sensorimotor performance asymmetries predict hand selection. Neuroscience 228:349–360 doi: 10.1016/j.neuroscience.2012.10.046

43. Rietdyk S, Drifmeyer JE (2010) The rough-terrain problem: accurate foot targeting as a function of visual information regarding target location. J Mot Behav 42:37–48 doi: 10.1080/00222890903303309

44. Rosenbaum D, Marchak F, Barnes H, Vanghan J, Slotta J (1990) Constraints on action selection: Overhand versus underhand grips. In: Jeannerod M (ed) Attention and performance XIII. Hillsdale, NJ: Lawrence Erlbaum Associates, pp 321–342

45. Rosenbaum DA, van Heugten CM, Caldwell GE (1996) From cognition to biomechanics and back: the end-state comfort effect and the middle-is-faster effect. Acta Psychol (Amst) 94:59–85 doi: 10.1016/0001-6918(95)00062-3

46. Schmidt RA, Lee T, Winstein C, Wulf G, Zelaznik H (2019) Motor Control and Learning. Human Kinetics

47. Schweighofer N, Xiao Y, Kim S, Yoshioka T, Gordon J, Osu R (2015) Effort, success, and nonuse determine arm choice. J Neurophysiol 114:551–559 doi: 10.1152/jn.00593.2014

48. Scott SH, Cluff T, Lowrey CR, Takei T (2015) Feedback control during voluntary motor actions. Curr Opin Neurobiol 33:85–94 doi: 10.1016/j.conb.2015.03.006

49. Shadmehr R, Huang HJ, Ahmed AA (2016) A Representation of Effort in Decision-Making and Motor Control. Curr Biol 26:1929–1934 doi: 10.1016/j.cub.2016.05.065

50. Shea JB, Morgan RL (1979) Contextual interference effects on the acquisition, retention, and transfer of a motor skill. Journal of Experimental psychology: Human Learning and memory 5:179

51. Smid KA, den Otter AR (2013) Why you need to look where you step for precise foot placement: the effects of gaze eccentricity on stepping errors. Gait Posture 38:242–246 doi: 10.1016/j.gaitpost.2012.11.019

52. Sparrow WA, van der Kamp J, Savelsbergh GJ, Tirosh O (2003) Foot-targeting in reaching and grasping. Gait Posture 18:60–68

53. Sterr A, O’Neill D, Dean PJ, Herron KA (2014) CI Therapy is Beneficial to Patients with Chronic Low-Functioning Hemiparesis after Stroke. Front Neurol 5:204 doi: 10.3389/fneur.2014.00204

54. Stewart JC, Gordon J, Winstein CJ (2013) Planning and adjustments for the control of reach extent in a virtual environment. Journal of NeuroEngineering and Rehabilitation 10:27 doi: 10.1186/1743-0003-10-27

55. Summerside E, Shadmehr R, Ahmed AA (2018) Vigor of reaching movements: reward discounts the cost of effort. J Neurophysiol doi: 10.1152/jn.00872.2017

56. Thomas MA, Fast A (2000) One step forward and two steps back: the dangers of walking backwards in therapy. Am J Phys Med Rehabil 79:459–461 doi: 10.1097/00002060-200009000-00011

57. Trommershäuser J, Maloney LT, Landy MS (2008) Decision making, movement planning and statistical decision theory. Trends Cogn Sci 12:291–297 doi: 10.1016/j.tics.2008.04.010

58. Wang J, Lum PS, Shadmehr R, Lee SW (2021) Perceived effort affects choice of limb and reaction time of movements. J Neurophysiol 125:63–73 doi: 10.1152/jn.00404.2020

59. Winter DA (1995) A.B.C. (anatomy, Biomechanics and Control) of Balance During Standing and Walking. Waterloo Biomechanics

60. Winter DA, Patla AE, Frank JS, Walt SE (1990) Biomechanical walking pattern changes in the fit and healthy elderly. Physical therapy 70:340–347

61. Winter DA, Pluck N, Yang JF (1989) Backward walking: a simple reversal of forward walking? J Mot Behav 21:291–305

62. Wolpert DM, Landy MS (2012) Motor control is decision-making. Curr Opin Neurobiol 22:996–1003 doi: 10.1016/j.conb.2012.05.003

63. Wu AR, Kuo AD (2016) Determinants of preferred ground clearance during swing phase of human walking. J Exp Biol 219:3106–3113 doi: 10.1242/jeb.137356

64. Wulf G, Lewthwaite R (2016) Optimizing performance through intrinsic motivation and attention for learning: The OPTIMAL theory of motor learning. Psychon Bull Rev 23:1382–1414 doi: 10.3758/s13423-015-0999-9

65. Yakovenko S, Drew T (2015) Similar Motor Cortical Control Mechanisms for Precise Limb Control during Reaching and Locomotion. J Neurosci 35:14476–14490 doi: 10.1523/JNEUROSCI.1908-15.2015

