## Supplementary Material for "Leg choice for volitional goal-directed stepping is primarily influenced by effort rather than success: A preliminary investigation in neurotypical adults"

**Supplementary Materials**

**Experiment 1**

**Methods**

*Experimental Procedures*

Given the overall aim of the study, we reduced the 12 target directions to 8 target regions. Four adjacent target directions (e.g., 1 and 2, 4 and 5, 7 and 8, 10 and 11) were consolidated except for four mid-horizontal and vertical target directions of 90/180/270/360 deg (i.e., 3, 6, 9, 12). This consolidation resulted in eight target regions (3 for the right hemispace, 3 for the left hemispace, and 2 for the central anterior-posterior axis) independent of target extent: Right-Anterior (RA; 1 and 2), Right-Medial (RM; 3), Right-Posterior (RP; 4 and 5), Central-Posterior (CP; 6), Left-Posterior (LP; 7 and 8), Left-Medial (LM; 9), Left-Anterior (LA; 10 and 11), Central-Anterior (CA; 12) (See **Fig. 1a**).

*Data Analyses*

Given that all participants were right leg dominant, we quantified the probability of choosing the right limb for the *Free* condition. For each participant, probability was calculated for each target, which was assigned three potential values: 0 – right leg was not chosen in either trial, 0.5 – right leg was chosen in one trial only, 1 – right leg was chosen in both trials. Then the average across participants was calculated for each target, which could potentially have a minimum value of 0 (i.e., no participant chose the right leg in either trial to step into this target) and a maximum value of 1 (i.e., all participants chose the right leg in both trials to step into this target).

The success of task performance was determined by calculating stepping error, which was used as a proxy for accuracy (i.e., the higher the stepping error the lower the accuracy). Stepping error was defined as a loss of balance, stepping on the wrong target, or missing the target. For each participant, stepping error was quantified for each target using the following rubric. There were three potential values: 0 – no stepping error in either trial, 0.5 – stepping error in one trial only, 1 – stepping error in both trials. Then the average across participants was calculated for each target which could potentially have a minimum value of 0 (i.e., no participant stepped into this target with error in either attempt) or a maximum value of 1 (i.e., all participants stepped into this target with an error in both attempts).

Responses from the self-perceived target difficulty survey were scored to determine which target(s) were perceived to be the most difficult. Self-perceived target difficulty was quantified by calculating the average across participants, where more than one target could be chosen. For each participant, perceived target-difficulty was calculated for each target, which was assigned two potential values: 0 – target was not selected to be difficult, 1 – target was selected to be difficult. Therefore, the potential minimum and maximum values per target was 0 and 1, respectively.

*Statistical Analyses*

| **Dependent Variables** | **Stepping into**  **ALL Target Regions during FREE** | | | |
| --- | --- | --- | --- | --- |
| Probability of Choosing the Right Leg | Two-way rmANOVA:  8 Target Region (RA, RM, RP, CP, LP, LM, LP, CA)  ×  3 Target Extent (Near, Middle, Far) | | | |
|  | **Stepping into the**  **LEFT Target Regions** | **Stepping into RIGHT Target Regions** | | **Stepping into the**  **CENTRAL Target Regions** |
| Stepping Error | Three-way rmANOVA:  2 Choice Condition (Free, Constrained)  **×**  3 Target Region (LP, LM, LA)  **×**  3 Target Extent (Near, Middle, Far) | Three-way rmANOVA:  2 Choice Condition (Free, Constrained)  **×**  3 Target Region (RP, RM, RA)  **×**  3 Target Extent (Near, Middle, Far) | | Three-way rmANOVA:  2 Choice Condition (Free, Constrained)  **×**  2 Target Region (CA, CP)  **×**  3 Target Extent (Near, Middle, Far) |
| Self-Perceived Target  Difficulty |  |  |  |  |
|  | **Non-Parametric associations (Spearman’s) between error and perceived difficulty per Choice condition** | | | |
| Stepping Error  &  Self-Perceived Target Difficulty | Free | | Constrained | |

**Table S1.** Statistical analyses (within-subjects design) used in Experiment 1.

rmANOVA: repeated measures Analysis of Variance; RA: Right-Anterior; RM: Right-Medial; RP: Right-Posterior; CP: Central-Posterior; LP: Left-Posterior; LM: Left-Medial; LA: Left-Anterior; CA: Central-Anterior.

**Experiment 2**

**Methods**

*Experimental Procedures:*

Modifications to the empirical approach used in Experiment 1: Due to observations and results of Experiment 1, we made the four following methodological modifications in Experiment 2. First, we eliminated both the central and medial targets. The elimination of the central targets (i.e., target directions: 6/CP/180 deg and 12/CA/360 deg; see **Fig. 1a**) was due to the limb choice inconsistency across participants to these targets (either leg was used with bias toward the right dominant leg). The elimination of the medial targets (i.e., target directions: 3/RM/90 deg and 9/LM/270 deg; see **Fig. 1a**) was because the stepping execution was very challenging across participants to these targets (i.e., target direction: 3/RM/90 deg) during *Constrained*. That left a total of 4 target regions ([left vs. right] × [anterior vs. posterior]) instead of the 8 used in Experiment 1. Second, we included a relevant set of kinematic measures to better characterize and quantify success (i.e., constant error and variable error) and effort (i.e., peak vertical foot lift and step path ratio); effort was not characterized in Experiment 1 but is integral to our research question. Including measures of effort in Experiment 2 provided a quantitative means to further explore the interactive effect of Choice (Free vs. Constrained) × Target Region ([Left vs. Right] × [Anterior vs. Posterior]) × Target Extent (Near vs. Middle vs. Far). Third, we increased the number of trials per target (i.e., from 2 to 3); this allows a determination of inter-trial variability, a more sensitive measure of skilled actions. Given that we reduced the number of targets and increased the number of attempts per target, the total number of trials per limb condition was the same as Experiment 1 (i.e., 72 trials). Fourth, we refined the assessment of self-perceived target difficulty to improve the quantification of subjective difficulty of a target by ensuring that all participants identify the most challenging target during each Choice condition.

Twenty-four targets (diameter: 0.05 m) were arranged in 8 target directions and classified into four target regions independent of target extent: Right-Anterior (RA), Right-Posterior (RP), Left-Posterior (LP), Left-Anterior (LA). In both Choice conditions, participants stood in a relaxed, comfortable stance at the central point of a center-out target array, facing a screen (1.5 m x 1.5 m), which was raised to each participant’s eye level and positioned 3 m in front of the wooden platform (1.82 m x 1.52 m x 0.03 m) on which instructions were displayed throughout the experiment using a projector. For both conditions, instructions were similar except for leg use. Participants were asked to prepare to step after a visual ‘ready’ and then ‘go’ signal that was synchronized with a simultaneous auditory signal (i.e., both visual and auditory cues were computer generated). The visual cue for each target lasted 6 seconds. Participants were instructed to aim and step with the ball of their foot onto the visually cued target as accurately as possible, keep their foot at the point of contact for about 1 second, and then return to the central position, completing a single uncorrected discrete movement. Throughout the test, participants were asked to keep their body facing forward towards the screen. When stepping to anterior targets (LA and RA), participants could use online visual input of the target, but not when stepping to posterior targets (LP and RP). The rest of experimental procedures were the same as in Experiment 1.

Each of the 24 targets was cued pseudo-randomly so that no target would be repeated successively. A full complement of trials consisted of three attempts to each target (24 x 3 = 72) per condition (a total of 144 trials per person). Targets were grouped in four blocks (6 targets per block); therefore, there were 12 blocks per condition. Data collection began 4 seconds before the onset of the first visual and auditory ‘go’ signal (beginning of a block) and ceased 4 seconds after the onset of the 6^th^ visual and auditory ‘go’ signal (end of a block). Data collection for each block took approximately 44 seconds. Between blocks, the participant was asked (via a visual cue) to stay in place during a 20-second break to prevent fatigue.

As in Experiment 1, augmented feedback was not provided in either condition whereas we assumed that all task-relevant intrinsic feedback (e.g., proprioceptive) was inherent to the conditions. During testing conditions, an experimenter visually monitored the participants to ensure compliance with the study protocol (e.g., visualizing anterior targets but not posterior, using the left leg during the *Constrained*, etc.)

*Data analyses*

The three-dimensional position data of the toe markers were analyzed offline using a custom written program in Matlab 9.10 (MathWorks Inc., Cambridge, MA, USA) software. Data were sampled at 125 Hz throughout each stepping trial and then digitally filtered using a low pass (10 Hz cut-off frequency), second order, and zero phase lag Butterworth filter. The temporal events of stepping (i.e., onset and offset) were calculated using standardized procedures used in our lab (Stewart et al. 2013).

After the signal processing, position-derived kinematic measures were calculated, first for each trial and then averaged across the per target trials. A total of six position-derived kinematic measures were calculated: constant error (mm), variable error (mm), peak vertical foot lift (mm), and step path ratio (unitless), foot path distance (mm), and movement distance (mm). The former four were used in statistical analyses as proxies of actual success (constant error and variable error) and motor effort (peak vertical foot lift, and step path ratio), whereas the latter two were used to calculate the step path ratio. Constant error was calculated as the linear distance between the marker position at movement offset (i.e., movement endpoint position) and the center of the target. Variable error was calculated as the square root of sum of squared differences between the movement offset (i.e., movement endpoint position) of each trial and the mean of the movement offset (x and y only) divided by the number of good trials (e.g., 3). The peak vertical foot lift was calculated as the highest vertical displacement of the second metatarsal during the stepping trial. Step path ratio was calculated as the ratio of the foot path distance (i.e., three-dimensional distance of the movement trajectory from movement starting point to movement end point) to movement distance (i.e., three-dimensional Euclidean distance from movement starting point to movement end point).

*Statistical Analyses*

**Table S2.** Statistical analyses (within-subjects design) used for the four kinematic measures in Experiment 2.

| **Dependent**  **Variable** | **Stepping into the**  **LEFT Target Regions** | **Stepping into the**  **RIGHT Target Regions** |
| --- | --- | --- |
| Constant  Error (mm) | Three-way rmANOVA:  2 Choice Condition (Free, Constrained)  **×**  2 Target Region (LA, LP)  **×**  3 Target Extent (Near, Middle, Far) | Three-way rmANOVA:  2 Choice Condition (Free, Constrained)  **×**  2 Target Region (RA, RP)  **×**  3 Target Extent (Near, Middle, Far) |
| Variable  Error (mm) |  |  |
| Peak Vertical  Foot Lift (mm) |  |  |
| Step Path  Ratio |  |  |

rmANOVA: repeated measures Analysis of Variance; LP: Left-Posterior; LA: Left-Anterior; RA: Right-Anterior; RP: Right-Posterior.

**References**

Stewart JC, Gordon J, Winstein CJ (2013) Planning and adjustments for the control of reach extent in a virtual environment. Journal of NeuroEngineering and Rehabilitation 10:27 doi: 10.1186/1743-0003-10-27

**Supplementary figures**

**1**

**2**

**11**

**10**

**4**

**5**

**7**

**8**

**Figure S1. Self-reported questionnaire for capturing self-perceived difficulty.** At the end of each Choice condition, each participant was asked to report: ***“****Overall, which target did you find the most challenging [_________]? (e.g. green 11)”*
